# The Medaka Inbred Kiyosu-Karlsruhe (MIKK) Panel

**DOI:** 10.1101/2021.05.17.444412

**Authors:** Tomas Fitzgerald, Ian Brettell, Adrien Leger, Nadeshda Wolf, Natalja Kusminski, Jack Monahan, Carl Barton, Cathrin Herder, Narendar Aadepu, Jakob Gierten, Clara Becker, Omar T. Hammouda, Eva Hasel, Colin Lischik, Katharina Lust, Risa Suzuki, Erika Tsingos, Tinatini Tavhelidse, Thomas Thumberger, Philip Watson, Bettina Welz, Nadia Khouja, Kiyoshi Naruse, Ewan Birney, Joachim Wittbrodt, Felix Loosli

## Abstract

Unraveling the relationship between genetic variation and phenotypic traits remains a fundamental challenge in biology. Mapping variants underlying complex traits while controlling for confounding environmental factors is often problematic. To address this, we have established a vertebrate genetic resource specifically to allow for robust genotype-to-phenotype investigations. The teleost medaka (*Oryzias latipes*) is an established genetic model system with a long history of genetic research and a high tolerance to inbreeding from the wild. Here we present the Medaka Inbred Kiyosu-Karlsruhe (MIKK) panel: the first near-isogenic panel of 80 inbred lines in a vertebrate model derived from a wild founder population. Inbred lines provide fixed genomes that are a prerequisite for the replication of studies, studies which vary both the genetics and environment in a controlled manner and functional testing. The MIKK panel will therefore enable phenotype-to-genotype association studies of complex genetic traits while allowing for careful control of interacting factors, with numerous applications in genetic research, human health, and drug development and fundamental biology. Here we present a detailed characterisation of the genetic variation across the MIKK panel, which provides a rich and unique genetic resource to the community by enabling large-scale experiments for mapping complex traits.

## Introduction

The relationship between natural genetic variation and the variance of quantitative traits in different species is one of the founding questions in genetics [1,2], and has become a very active field of research today. Experimentally it has been addressed using plant and animal models [3–5], and has been studied in human populations [6–8]. In model organisms such as *Arabidopsis thaliana* and *Drosophila melanogaster*, genome-wide association studies (GWAS) and specific crosses have been used to examine complex genetic traits, bridging population association models to more traditional controlled-cross strategies [9,10]. GWAS have had tremendous success in discovering genomic loci underlying human traits by leveraging observational outbred cohorts of individuals [11]. However, this outbred sampling strategy still leaves many questions in vertebrate genetics unanswered, including the importance of gene-by-environment interactions (GxE) and epistatic interactions, i.e. between two genetic loci (GxG) – for phenotypic extremes in particular – as well as the potential combination of both interactions (GxGxE) [12,13]. To explore these questions, one must turn to laboratory vertebrates where one can vary and control both genetic and environmental sources of variance.

The mouse (*Mus musculus*) is an established vertebrate model for human genetics, and researchers have created panels of recombinant inbred lines (RILs), such as the Collaborative Cross and the BXD cross [14,15], in order to run GWAS in these managed populations [16] and have a controlled source of genetic variation. As mice are mammals, they have excellent orthologous organ systems and cell types to humans, and an unsurpassed repertoire of tools to control their precise genome, including large scale genomic engineering [17,18]. However, the genetic variation of all laboratory mice follows from the complicated history of the domestication of “fancy mice”, originating from three separate species that were bred in captivity, and then undergoing complex domestication before the laboratory lines were established [19,20]. As such, “polymorphisms” between laboratory strains and GxG (epistatic) interactions result from the non-natural creation of these lines. Furthermore, even large panels of RILs, such as the Collaborative Cross and BXD, come from a limited number of parents [21]. The unnatural genetic origin of laboratory mice and their limited parentage means that they have deficiencies in modelling outbred populations such as humans. It would be optimal to supplement the mouse RIL lines with other vertebrate species both to better capture outbred settings, and to provide another window into vertebrate genetics that can be controlled in the laboratory. In addition, association studies often require high numbers of phenotyping experiments, so the ease of phenotyping and economical husbandry are also important features for a suitable vertebrate animal model.

The teleost medaka (*Oryzias latipes*) is a long-established model organism that combines a number of important features [22]. Economical husbandry, high fecundity, and transparency of embryos and largely also adults, allow one to carry out a wide range of phenotyping on large numbers of individuals. Medaka has a long history in genetic research [2,23] and comes with a wide range of established molecular genetic tools that allow in-depth analysis of gene function and phenotypes [24], including CRISPR/Cas-based homologous recombination protocols that permit the precise testing of genetic variants in different inbred strains [25]. Over 70% of human genes have teleost orthologs, and nearly all the major organ systems in humans have a teleost counterpart [26,27], making it possible to translate many findings between species.

Importantly, medaka is highly tolerant to inbreeding from the wild [28]. Isogenic inbred medaka strains have been established from natural populations, and a number of these laboratory strains have been inbred for over a hundred generations [29]. This indicates that medaka’s tolerance to inbreeding is a species-specific trait, rather than depending on the starting population. These medaka isogenic inbred strains have been bred by single full-sibling-pair (brother-sister) crosses [30]. Such full-sibling-pair crosses are commonly used for inbreeding in medaka as they confer a strong genetic bottleneck in each generation without requiring the maintenance of more than one generation per cross, as opposed to “backcrosses” which involve crossing F1 individuals with their parents.

We have previously identified a polymorphic wild population, sampled from Kiyosu, Japan [31]. By single full-sibling-pair inbreeding for 9 generations, we have established a panel of 80 near-isogenic inbred lines from this wild population, known as the Medaka Inbred Kiyosu-Karlsruhe (MIKK) panel. In this paper we describe this panel, outline the success of this inbreeding strategy, and provide a view of the lines based on their whole-genome sequences. We show that both molecular and organismal phenotypes are distinguishable in this panel and that molecular traits can be mapped to specific loci. We also comment on aspects of medaka genetics and genomic biology from the deep sequencing of the MIKK panel, from population genetics to loss of function alleles and structural variation. We show that the MIKK panel is an inbred near-isogenic resource with a good representation of wild alleles in the Kiyosu population, and is well suited for future genetic association and mapping studies.

## Results

### Inbreeding of the MIKK panel

We have previously reported the identification of a wild medaka population that satisfies critical conditions for the establishment of a vertebrate near-isogenic panel [31]. In July 2010, we selected a genetically diverse medaka population from the Kiyosu area near Toyohashi, Aichi Prefecture, and determined that it was free of significant population structure and introgression from aquarium populations. Segregation analysis of trios revealed advantageous linkage disequilibrium (LD) properties with LD estimates expressed as *r*^2^ reaching a minimum of ~0.12 at a distance of approximately 12.5 kb.

To commence the inbreeding process, we first established founder families by setting up 115 random crosses of single mating pairs from the wild Kiyosu population. For each founder family, we then set up mostly two, but up to five, single full-sibling-pair inbreeding crosses. This resulted in a total of 253 single full-sibling-pair crosses for the F1 generation, which we used to initiate the inbreeding lines. Lines derived from the same founder family are referred to as ‘sibling lines’. In the subsequent 8 generations of inbreeding, we only used one mating pair per line. 19 of the F1 crosses did not result in productive mating, so we proceeded to inbreed the F2 generation with 234 productive single full-sibling-pair crosses.

During the first 9 generations of inbreeding (using only one mating pair per line), to avoid selecting for specific traits, we intentionally did not consider body size, generation time, fecundity, fertilization rate, survival rate or sex ratio. We continued inbreeding with all lines independent of these traits up to the 9th generation, or until the strain became extinct (**Figure 1A**). The only criterion we considered when selecting individuals for inbreeding crosses was average wild-type morphology (irrespective of size), so we did not use individuals with severe malformations, such as a strongly-bent tail. When all fish in a given line showed the same morphological abnormalities, for example line 4-1 which has an unusually large abdomen, we continued inbreeding with these fish, assuming a genetic cause for the malformation.

**Figure 1:**
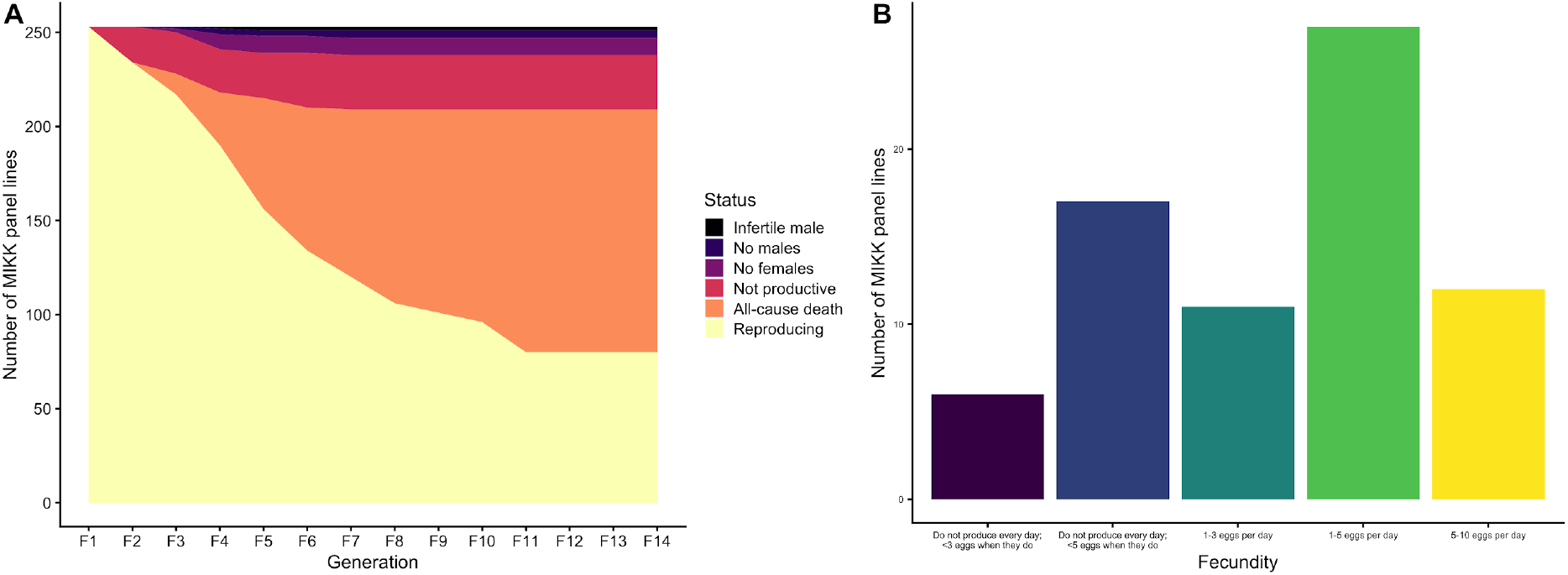
Inbreeding and fecundity of the MIKK panel lines. A: Status of all MIKK panel lines during the first 14 generations of inbreeding, showing cause of death for non-extant lines. B: Average fecundity of MIKK panel lines in generation F16, as measured during peak egg production in July 2020.

The most frequent cause for extinction of a line during inbreeding was all-cause mortality (**Figure 1A**). In the majority of cases, individuals died during the first 3 weeks after hatching due to unknown causes. The highest incidence of line extinction occurred while inbreeding generations 3 to 5. Dissection of dead fish revealed a severe infection of inner organs with *Flavobacterium columnare.* Antibiotic treatment stopped mortality within 24 hours, indicating that the bacterial infection was the main cause of death. Outbred Kiyosu fish were not affected; we therefore assume that inbreeding led to a higher susceptibility to bacterial infections. In addition, 13 lines were lost due to extreme shifts in the sex ratio. We also observed unproductive crosses in 10 lines, and we terminated the lines when 4 alternative sibling crosses from the line were all unproductive. At the time of submission, the MIKK panel generation F18 comprised a total of 80 near-isogenic inbred lines (column B, **Supplementary Table 1**).

### Fecundity and morphological measures across the MIKK Panel

We assessed fecundity of the MIKK lines at generation F16. Over a period of 4 weeks, we monitored each line’s egg production and assigned them to different classes of fecundity based on the approximate number of eggs produced per day. This assignment was facilitated by the fact that fertilized eggs remain attached to the females after mating. Under the constant summer conditions of the fish facility (14 hours light/10 hours dark), medaka mate every day [32]. This made it possible to semi-quantitatively measure fecundity by visually inspecting the number of females with fertilized eggs, and estimating the number of eggs per female that had mated. We observed a spectrum of mating frequencies and eggs per mating that ranged from occasional, irregular mating with few eggs per female, to daily mating with up to 10 eggs per female. For the majority of lines we observed daily mating with 1-3 or more eggs per female (**Figure 1B** and **Supplementary Table 2**). Thus the overall fecundity of the majority of MIKK lines is sufficient to meet the demands for phenotyping quantitative traits.

To assess some basic morphological differences across the MIKK panel, we took high resolution images of 77 lines for one male and two female fish aged between 6 and 9 months post-hatching, imaged in both lateral and dorsal orientations (**Supplementary File 1**). This resulted in 462 images which we used to assess whole-body anatomical differences and heritability across the MIKK panel. We applied image segmentation using modern machine learning techniques (**Methods**) allowing the extraction of specific morphometric parameters. We then used these parameters to calculate heritability estimates across the MIKK panel. To exclude confounding factors such as differences in developmental stage, we only extracted information that would allow us to generate relative measurements within each fish. For most measurements we used the overall number of pixels within the entire segmented body shape as the quantitative within-fish normalisation value.

We focused on three specific measurements to assess phenotypic differences: 1) the relative size of the eye from lateral view images; 2) the relative distance between the eyes (a measure of head size) from the dorsal view images; and 3) abdominal size in females from the lateral view. To measure the relative eye size across MIKK panel lines, we used the total number of pixels within the segmented eyes divided by the total length from nose to tail (in pixels), which produced a relative measure that corrects for the variance of body size. We performed one way analysis of variance (ANOVA) tests, calculating effect size (*η*^2^) as a measure of broad sense heritability (*H*^2^). The amount of variance explained by MIKK panel lines overall, and thus broad sense heritability (*H^2^*) is 0.68. We performed similar analyses for the relative distance between eyes and abdominal size parameters, observing 0.51 and 0.39 levels of heritability for these traits respectively (**Supplementary Table 3**).

Overall, the between-line variance for these simple morphological measurements across the MIKK panel is high compared to the within-line variance, and broadly similar to human and mouse morphometric measurements, suggesting that these traits are heritable in medaka as expected.

### Homozygosity of the MIKK panel genomes

We extracted DNA from dissected whole brains and performed whole genome sequencing across the panel, with per-sample coverage between 21-34x. As expected, over 97.3% of the reads aligned to the *HdrR* reference genome, and we called single nucleotide polymorphisms (SNPs) to this reference using a standard pipeline (**Methods**). We calculated the level of homozygosity across the medaka genome for each MIKK panel line using the number of heterozygous SNP calls across the genome. The medaka sex chromosome, LG1, was excluded in this heterozygosity estimation since heterogametic XY males were used for whole genome sequencing. However, as expected, this heterozygosity is not distributed evenly across the genome in each line. To explore this we split the genome up into 10 kilobase (kb) non-overlapping windows and counted the number of heterozygous SNPs in each window. We then trained a two-state Hidden Markov Model (HMM) on all 10 kb windows from all MIKK panel lines jointly, using the expectation maximisation (EM) algorithm to separate out true homozygous regions from regions of residual heterozygosity (**Methods**). We then used the Viterbi algorithm to find the most likely sequence of hidden states (Viterbi path) for each panel strain separately. We performed a similar procedure on the classic inbred strains (*iCab* and *HO5*) and wild-caught medaka from the same founding population as the MIKK panel [31]. Below are summary plots showing the proportion of called homozygous regions comparing the MIKK panel to classical inbred strains and wild Kiyosu medaka.

The genome-wide homozygosity observed in the MIKK panel lines is similar to that of the classical laboratory inbred strains (*iCab* and *HO5*) which have been inbred for more than 70 generations. Both classical inbred strains and the MIKK panel show a clear difference in the number of homozygous regions genome-wide relative to wild Kiyosu samples (**Figure 2A**). Over 70% of MIKK panel lines are homozygous for greater than 70% of their entire genome, which is equivalent to the similar inbred *Drosophila* resources [5]. As expected, when looking at the level of homozgosity across chromosomes, we observe a clear decrease on chromosome 1 across all MIKK panel lines (**Figure 2B**), reflecting that the sex-determination region must nearly always remain heterozygous in male medaka [33]. In addition, chromosomes 2 and 3 showed higher levels of heterozygosity (see discussion) compared to the other chromosomes. At the time of sequencing, all MIKK panel lines had undergone 9 generations of single full-sibling-pair inbreeding, and although the majority of lines have highly homozygous genomes, there is a subset of lines that show a considerably lower degree of homozygosity (**Figure 2C**). Overall, the MIKK panel shows similar levels of homozygosity compared to classical laboratory inbred medaka strains, and contains a strong increase in isogenic genotypes compared to wild medaka from the original wild population.

**Figure 2:**
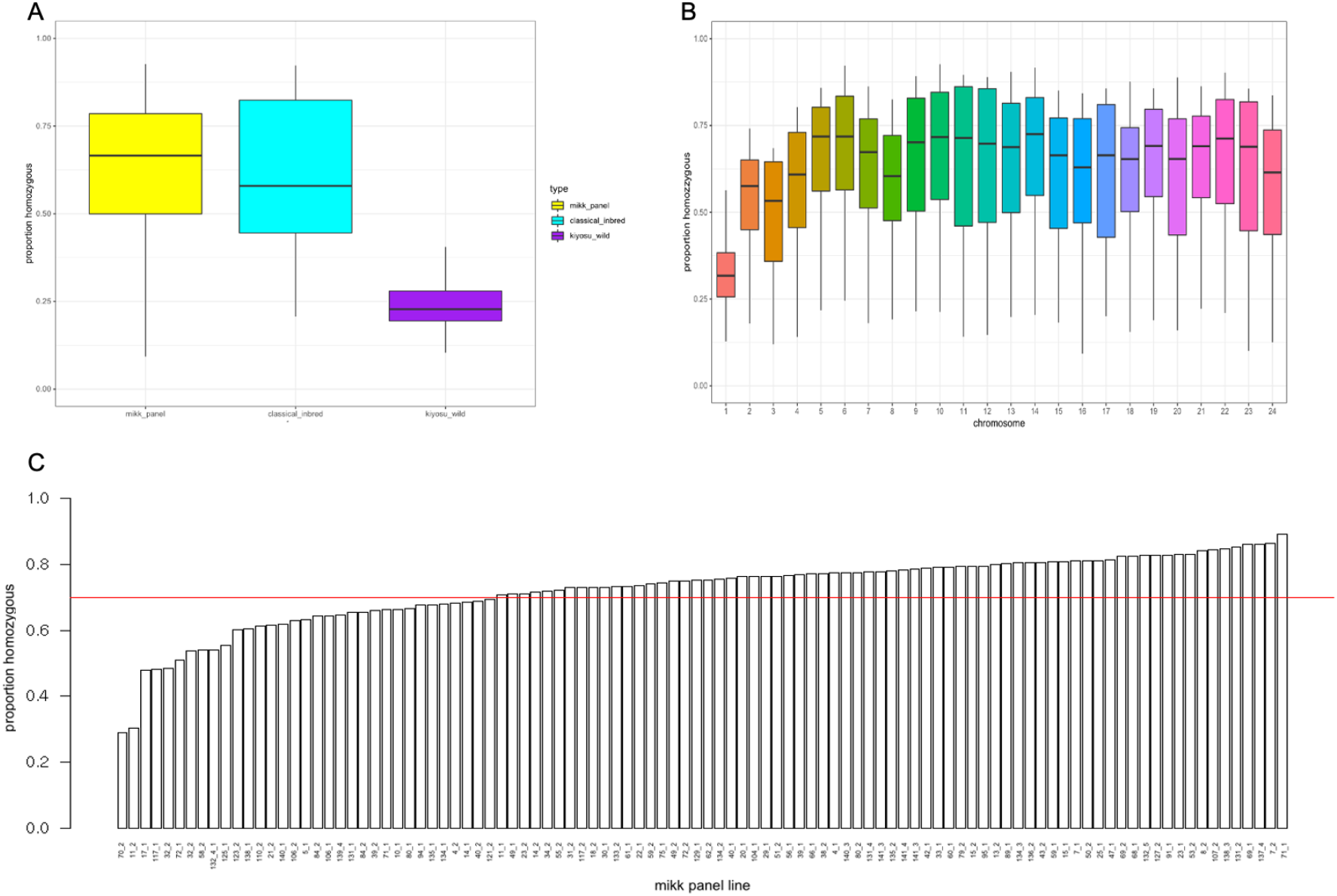
Homozygosity of the MIKK panel. **A**: The overall proportion of homozygous 10 kb genomic regions comparing the MIKK panel to classical inbred and wild Kiyosu fish. **B**: The proportion of homozygous 10 kb genomic windows across the MIKK panel for each chromosome. **C**: The proportion of homozygous 10 kb genomic windows (excluding chr1) for all MIKK panel lines.

### Enhanced definition of the medaka sex-determination region

To further investigate the homozygosity levels on chromosome 1, we used a 4-state HMM trained on the number of heterozygous SNPs in 10 kb windows. This created a finer-grained view across all MIKK panel lines, allowing us to estimate the critical region containing important genes necessary for sex determination in medaka [34].

The sequenced male XY regions on chromosome 1 that are maintained in their heterozygous state across MIKK panel lines refine the critical region for sex determination in medaka (**Figure 3A**). We determined the critical region using two different approaches. First, for all 10 kb windows, we calculated the proportion of lines of the MIKK panel that had greater than 5 heterozygous SNPs (fitted blue line in **Figure 3B**). Second, we used the HMM states to create a weighted estimate where states with higher heterozygosity levels contributed more weight in the model (fitted red line in **Figure 3B**). Overall, when using the SNP-count-based estimate, we observe a large region across the middle of chromosome 1 that remains heterozygous in most MIKK panel lines. However, when incorporating further information from the HMM, we detected a 0.5 Megabase (Mb) region (chr1:15959156-16459156) that showed considerably higher levels of heterozygosity in most MIKK panel lines. This region includes 17 genes in the *HdrR* reference genome and could be of interest for further mapping studies to investigate the precise location of the duplicated copy of the *DMRT1* gene known to be critical in sex determination [35], and how the suppression of recombination creates hetreogametic sex determination. We note that a small number of lines were homozygous across this region, consistent with occasional XX male sex determination seen in medaka (see discussion).

**Figure 3:**
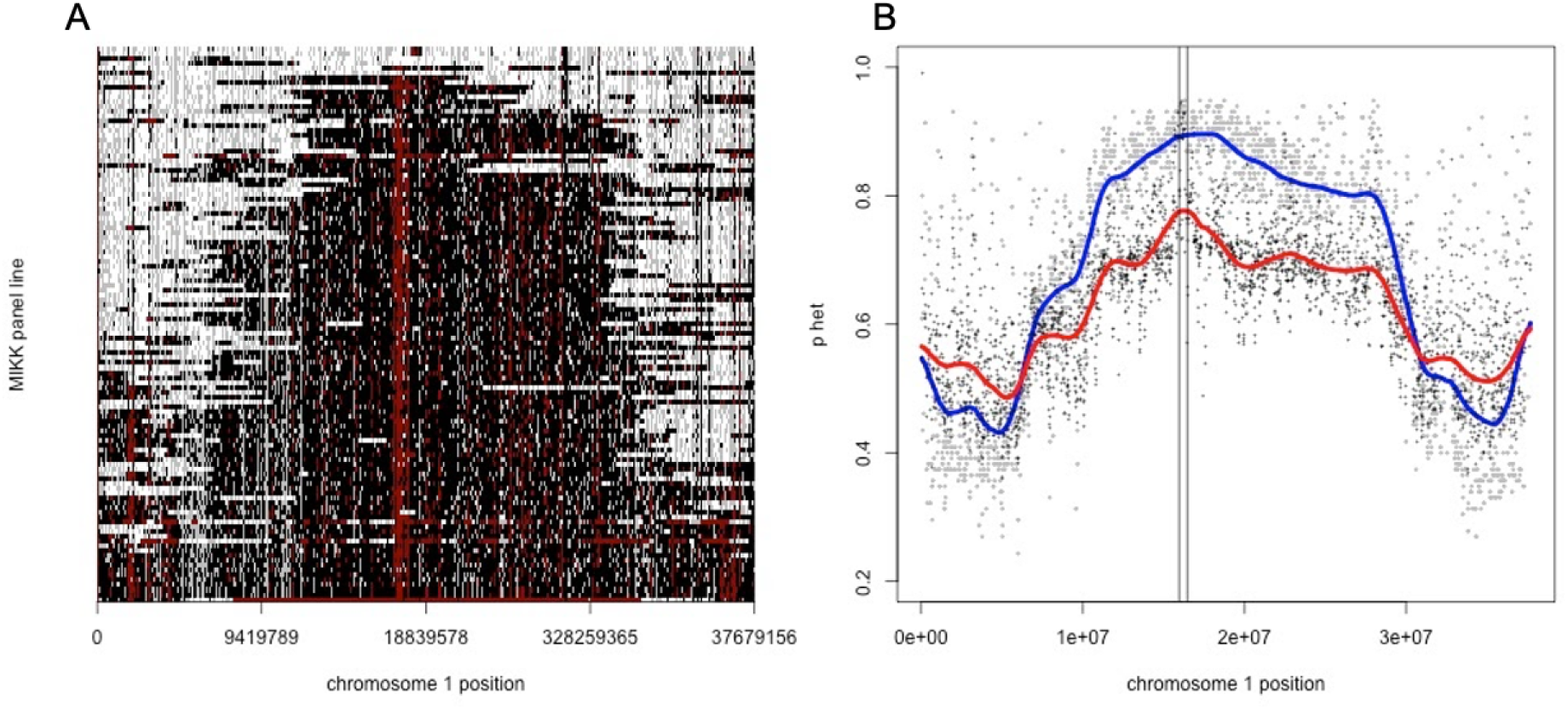
Heterozygosity across chromosome 1 in the MIKK panel defines the sex determining region. **A**: Homozygosity state from the HMM for all 10 kb windows across all MIKK panel lines (the colour ordering white, grey, black, and red represent increasing levels of heterozygosity). **B**: Two different estimates of the proportion of MIKK panel lines that are classified as heterozygous across all 10 kb windows on chromosome 1 (blue: based on heterozygous SNP count; red: based on HMM state space).

### Potential functional impact of small genomic variation

We discovered a total of 3,001,493 variants (SNPs and INDELs) across the MIKK panel compared to the *HdrR* reference, of which more that 70% (2,248,228) were either synonymous or intergenic (**Supplementary Table 4**). We examined the potential functional impact of variation in protein-coding genes across the panel. There were a total of 644,509 non-synonymous variants, 36,444 splice site variants and 82,312 potential Loss-of-Function (LoF) variants (stop codons or frameshifts). Similar to human studies [36], apparent LoF variation is enriched in regions where the genome assembly or gene predictions were poor, so we used a variety of computational screens (**Methods**) to select 35,154 high-confidence LoF variants in the MIKK panel (**Figure 4**).

**Figure 4:**
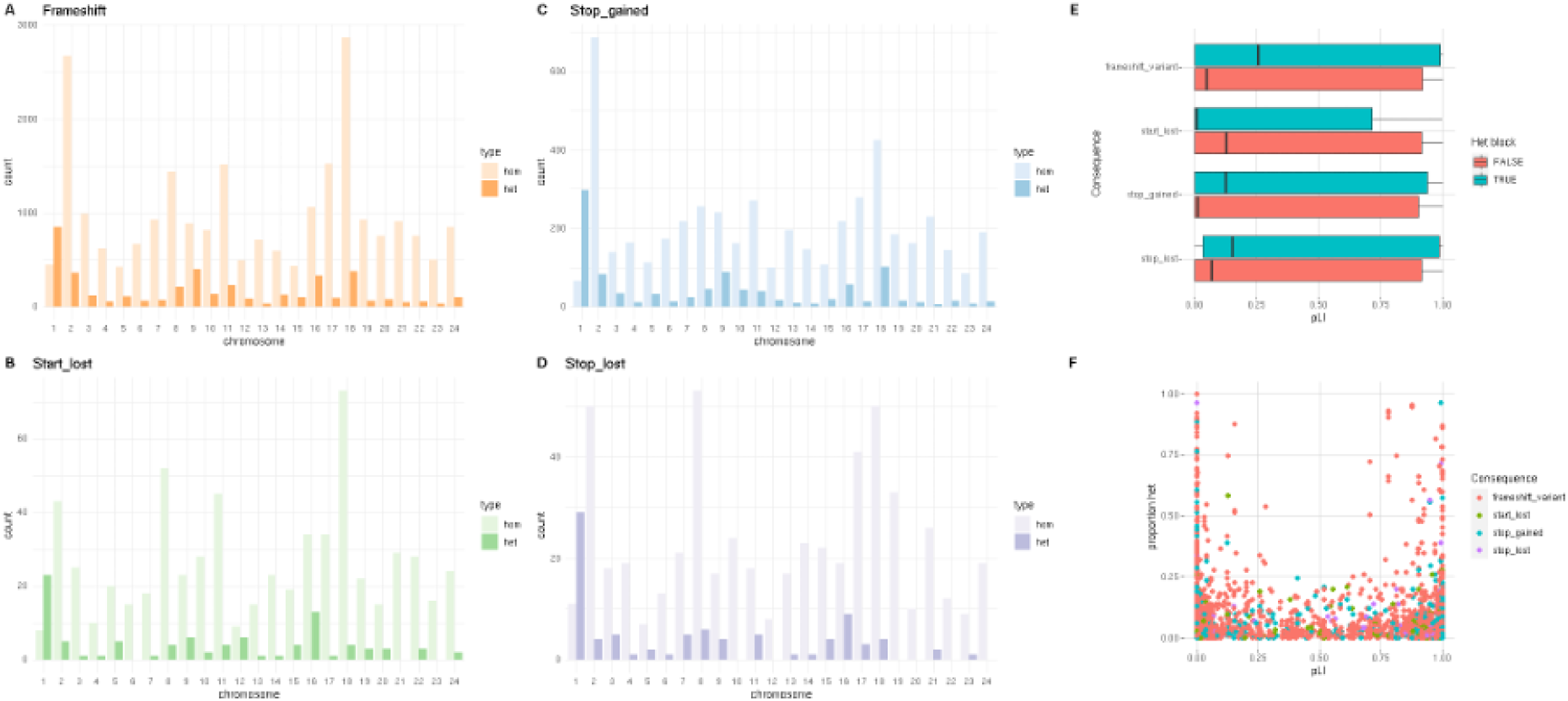
High-confidence loss-of-function variants in the MIKK panel. **A**: Frameshift mutations per chromosome within and outside heterozygous blocks. **B**: Start-lost mutations per chromosome within and outside heterozygous blocks. **C**: Stop-gained mutations per chromosome in and outside heterozygous blocks. **D**: Stop-lost mutations per chromosome within and outside heterozygous blocks. **E**: Variant class against purifying selection (pLI) within and outside heterozygous blocks. **F**:Purifying selection (pLI) against the proportion of genotypes in MIKK panel lines that are heterozygous.

As expected, these high-confidence LoF variants were more likely to be found in a heterozygous state (15% of the LoF SNPs were present in a heterozygous-called block compared to 9% on average), consistent with successful homozygosity being prevented in some cases due to a deleterious LoF allele. However, 63% had confident homozygous calls, with 34% of LoF SNPs showing no evidence of heterozygosity in any of the individual lines (**Figure 4F**). Unsurprisingly, we see the expected increase in the number of LoF variants in heterozygous blocks for chromosome 1 due the sex determination region, and interestingly both chromosomes 2 and 18 show consistently higher numbers of LoF variation for all LoF variant classes (**Figure 4A-D**) (see discussion).

Previous work in humans has characterised genes for their intolerance to mutations using the probability of LoF intolerance (pLI scores) which has been widely adopted in human studies [37]. For most LoF variant classes there is a slight enrichment towards higher pLI scores of the orthologous human genes inside heterozygous blocks compared to homozygous blocks (**Figure 4E**). This high-confidence, homozygous set of LoF variants were less present in medaka genes orthologous to human genes under stringent purifying selection, or disease-causing genes in humans (**Figure 4E,F**). There were 1,441 cases with a homozygous high-confidence LoF in the MIKK panel orthologous to a human dosage-sensitive gene (pLi > 0.5), listed in **Supplementary Table 5**. These individual variants, and the lines that carry them, may be of direct interest to researchers focused on these critical human genes.

### Identity-by-descent (IBD) mapping

To assess the population structure of the MIKK panel, we performed an identity-by-descent (IBD) analysis across the panel, including both classical inbred strains (*iCab* and *HO5*) and MIKK panel lines. As in previous studies, overall IBD is tracked via the identity-by-state (IBS) of SNPs. We first sought to assess the overall relationship between the classical inbred strains and the MIKK panel using IBS measures (**Figure 5A**). We observe a clear separation between the classical inbred strains and MIKK lines, reflecting comparably low recent shared ancestry.

**Figure 5:**
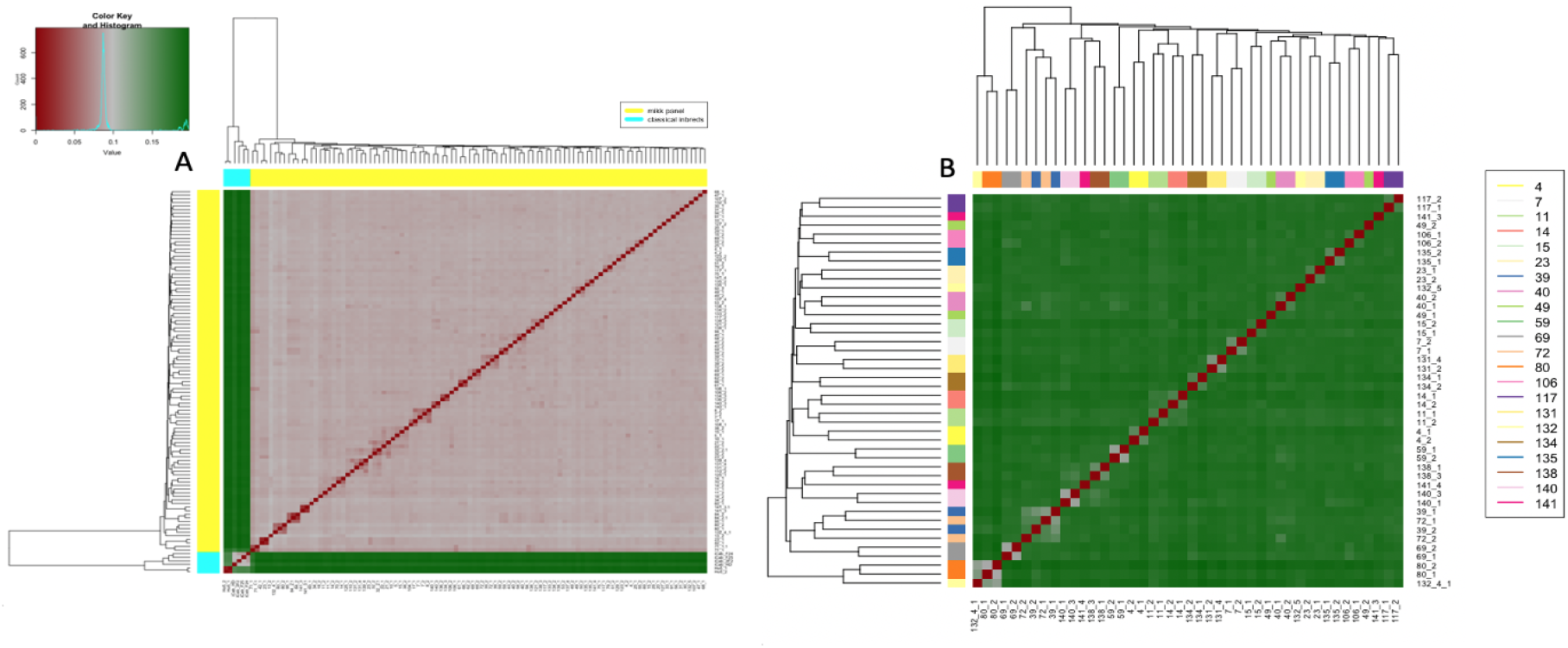
Identity by state (IBS) across the MIKK panel and classical inbred strains. **A**: 1-IBS from all medaka samples, including classical inbred strains. **B**: 1-IBS for all MIKK panel lines that have an available “sibling line”.

Next we compared IBS measures for all MIKK panel lines that had a ‘sibling line’ available (as described above). Therefore these sibling lines can be expected to share 25% of the genome from their parents. At the time of sequencing, there were 44 MIKK panel lines with a remaining productive sibling line (22 sibling-line pairs) (**Supplementary Table 1**). When we cluster sibling-line pairs based on IBD (**Figure 5B**), 77% (17 pairs) are directly adjacent, showing a higher degree of relatedness within the sibling pair than without. Two sibling-line pairs cluster as a quartet, suggesting that the founding Kiyosu individuals for these two pairs may have been closely related. As expected, we find that the majority of sibling-line pairs show a high degree of relatedness, which not only provides interesting population structure across the panel, but also shows that the careful and detailed tracking of the MIKK panel lines throughout the inbreeding process has been robust. The three “orphan” sibling line pairs which do not cluster directly adjacent to each other may well have arisen from random segregation in the inbreeding producing more divergent sib lines.

### Genetic composition of the MIKK panel

The MIKK panel was created from a single southern-Japanese population of wild medaka from Kiyosu, Japan. We sought to determine whether there had been any large-scale selection or allele skew on panel genotypes from the original wild founding population during the inbreeding process. To search for allele skew, we used the parents from eight medaka trio datasets collected from the same founding population [31] to measure population differentiation due to genetic structure with the fixation index (*F*_*ST*_). *F*_*ST*_ is a well-established parameter of population differentiation [38]. It provides a measure of genetic structure based on the variance of allele frequencies or allelic skew between populations [39]; high *F*_*ST*_ indicates that allele distributions between populations are divergent, whereas low *F*_*ST*_ indicates they are similar.

We calculated *F*_*ST*_ across all shared genotype positions between 16 wild medaka sequences [31] and the MIKK panel. To look at *F*_*ST*_ genome-wide, we defined 10 kb non-overlapping windows and took the mean *F*_*ST*_ for each window across the medaka genome (**Figure 6A**). Overall we see very little allele skew genome-wide, with no large regions showing high *F*_*ST*_ values. When looking at the 10 kb windows that display higher *F*_*ST*_ values, we see that the majority were based on windows with low numbers of available SNPs (**Figure 6C**). Over 93% of 10-kb windows had greater than 50 SNPs available to include when calculating the mean *F*_*ST*_, with a median of 115 sites per window (**Figure 6B**). Reassuringly, we did not observe any significant differences in the total genetic variance genome-wide between the MIKK panel and wild medaka samples from the same founding population. This shows that the genetic diversity within the MIKK panel is a reasonable representation of the genetic diversity found in wild medaka from the Kiyosu region.

**Figure 6:**
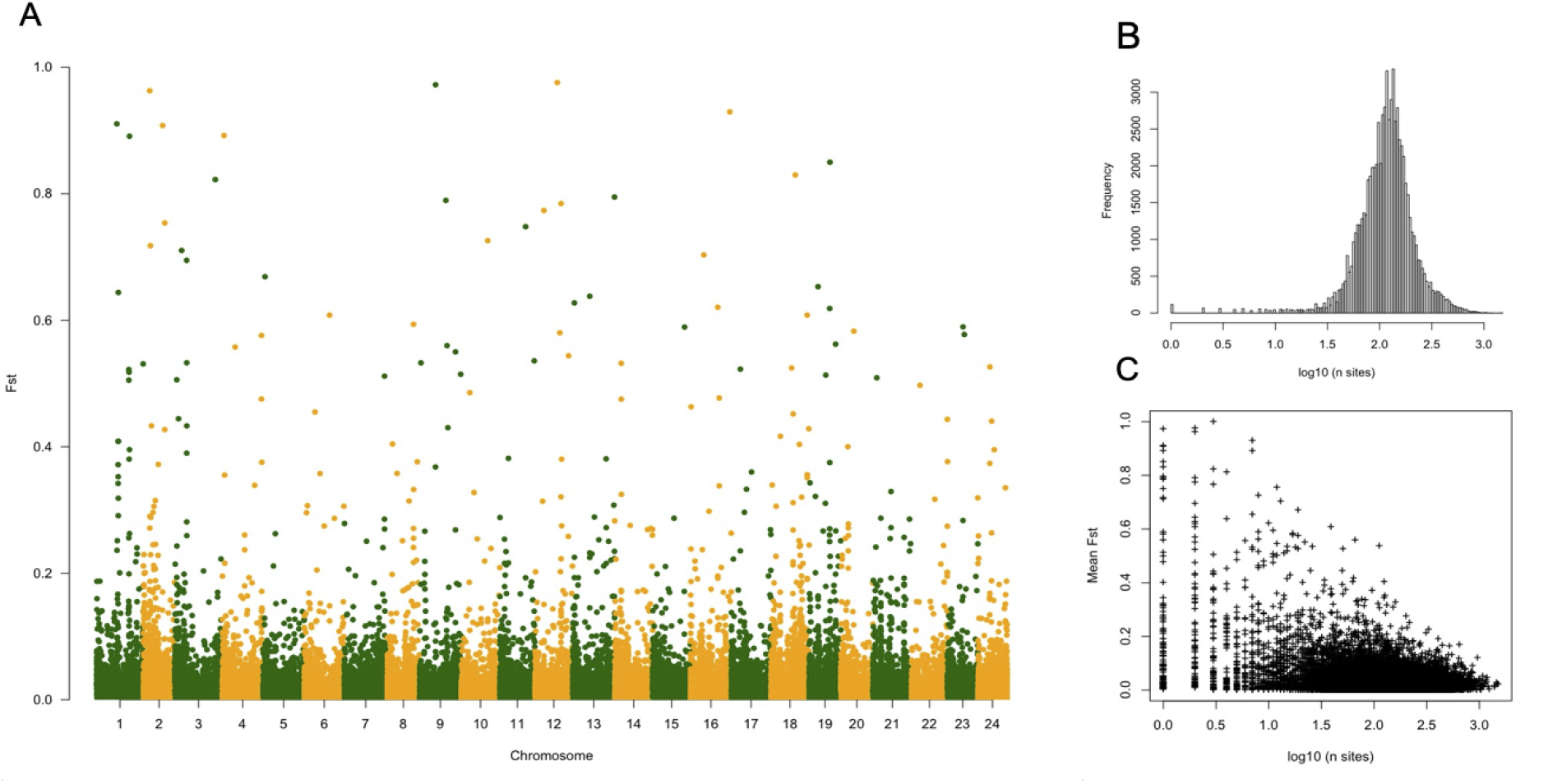
Fixation Index (F_ST_) comparing allele frequencies of the MIKK panel against wild Kiyosu medaka. **A**: Mean F_ST_ in 10-kb windows across the medaka genome. **B**: log_10_ of the number of SNPs per 10-kb window. **C**: log_10_ of number of SNPs per window vs mean F_ST_.

### Introgression with northern Japanese and Korean medaka populations

To determine whether the MIKK panel’s founder population showed signs of introgression with northern Japanese and Korean medaka populations, we ran an ABBA-BABA analysis [40–42]. We used the 50-fish multiple alignment in Ensembl’s release 102 [43] to obtain the aligned genome sequences of the established inbred medaka strains *HdrR* (southern Japan), *HNI* (northern Japan), and *HSOK* (Korea), in order to orientate each SNP to its ancestral state (**Figure 7A**). We then combined this data with the Illumina-based SNP calls for the MIKK panel and the established inbred medaka strain *iCab* (southern Japan). We used this combined dataset to calculate 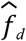 *[42]*, a modified version of the ‘admixture proportion’ statistic 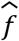 [40], to measure the proportion of shared genome in 500-kb sliding windows between the MIKK panel and either *iCab*, *HNI*, or *HSOK*. For this purpose we used *HdrR* as the MIKK panel’s most closely-related P1 population, the MIKK panel as the P2 focal population, either *iCab*, *HNI*, or *HSOK* as the P3 introgressing population, and the most recent common ancestor as the outgroup (O) (**Figure 7B**). 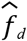 measures across the medaka genome are presented in **Figure 7C**. Based on the genome-wide mean 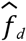, the MIKK panel shares approximately 25% of its genome with *iCab*, 9% with *HNI*, and 12% with *HSOK*. These results provide evidence that the MIKK panel’s originating population has more recently introgressed with medaka from Korea than with medaka from northern Japan. This supports the findings in [31], where the authors found little evidence of significant interbreeding between southern and northern Japanese medaka since the populations diverged.

**Figure 7:**
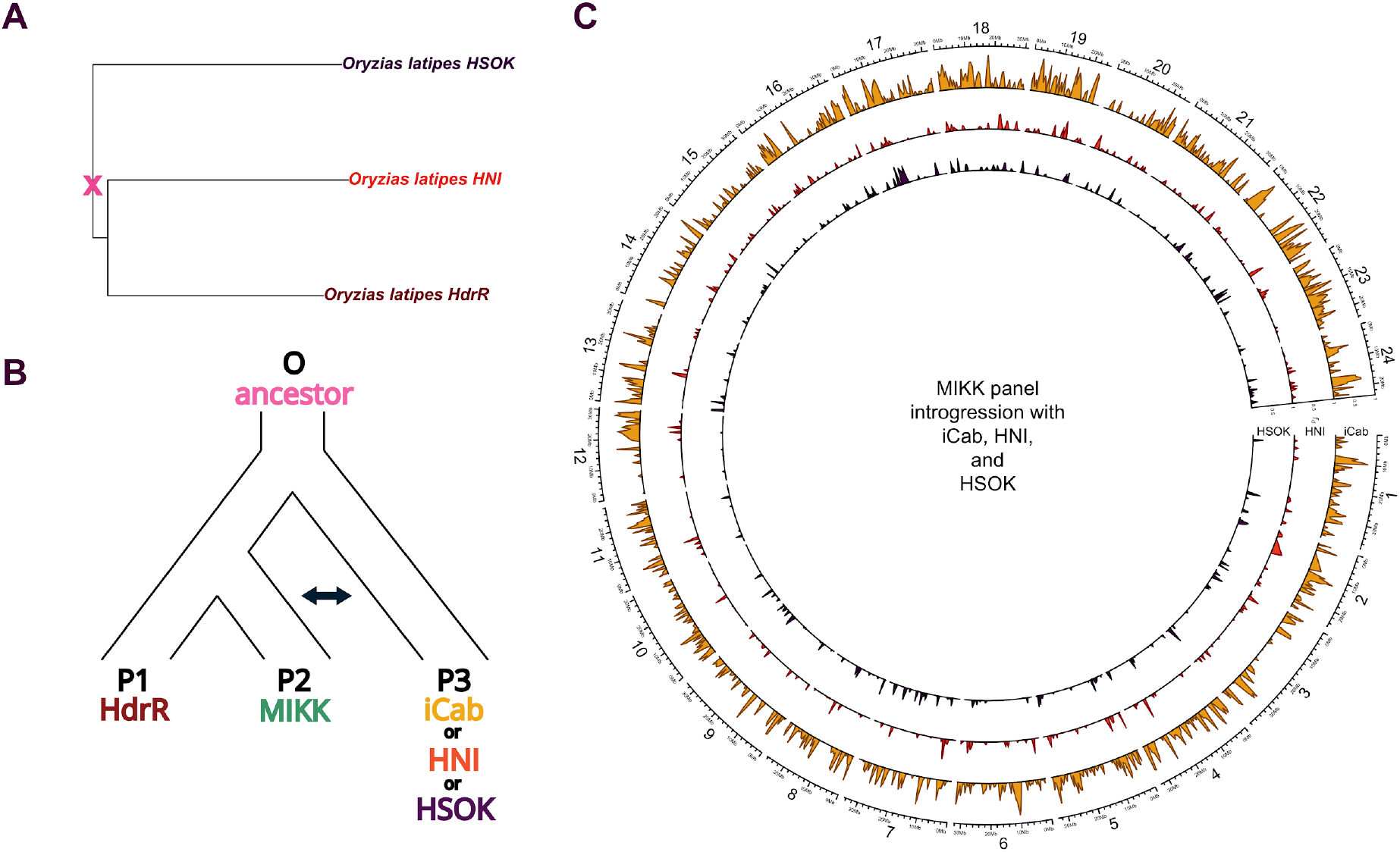
ABBA-BABA analysis. **A**. Phylogenetic tree generated from the Ensembl release 102 50-fish multiple alignment, showing only the medaka lines used in the ABBA-BABA analysis. **B**. Schema of the comparisons carried out in the ABBA-BABA analysis. **C**. Circos plot comparing introgression 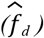 between the MIKK panel and either iCab (yellow), HNI (orange), or HSOK (purple), calculated within 500-kb sliding windows using a minimum of 250 SNPs per window.

### LD decay

We analysed the MIKK panel’s allele frequency distribution and linkage disequilibrium (LD) structure to assess their likely effects on genetic mapping. To remove allele-frequency biases introduced by the presence of “sibling lines” in the MIKK panel, we first filtered the Illumina-based variant calls to include only one inbred line from each pair, leaving *N* = 63 non-sibling inbred lines. **Figure 8A** compares the allele frequency distribution for the 16.4M biallelic SNPs identified in those filtered calls, with the allele frequency distribution of the 81M biallelic SNPs in the 1000 Genomes Project Phase 3 release (*N* = 2,504) (1KG) in human [44]. As expected, the 1KG and MIKK panel calls are similarly enriched for low-frequency variants, albeit to a lesser extent in the MIKK panel, which is likely due to its smaller sample size.

**Figure 8:**
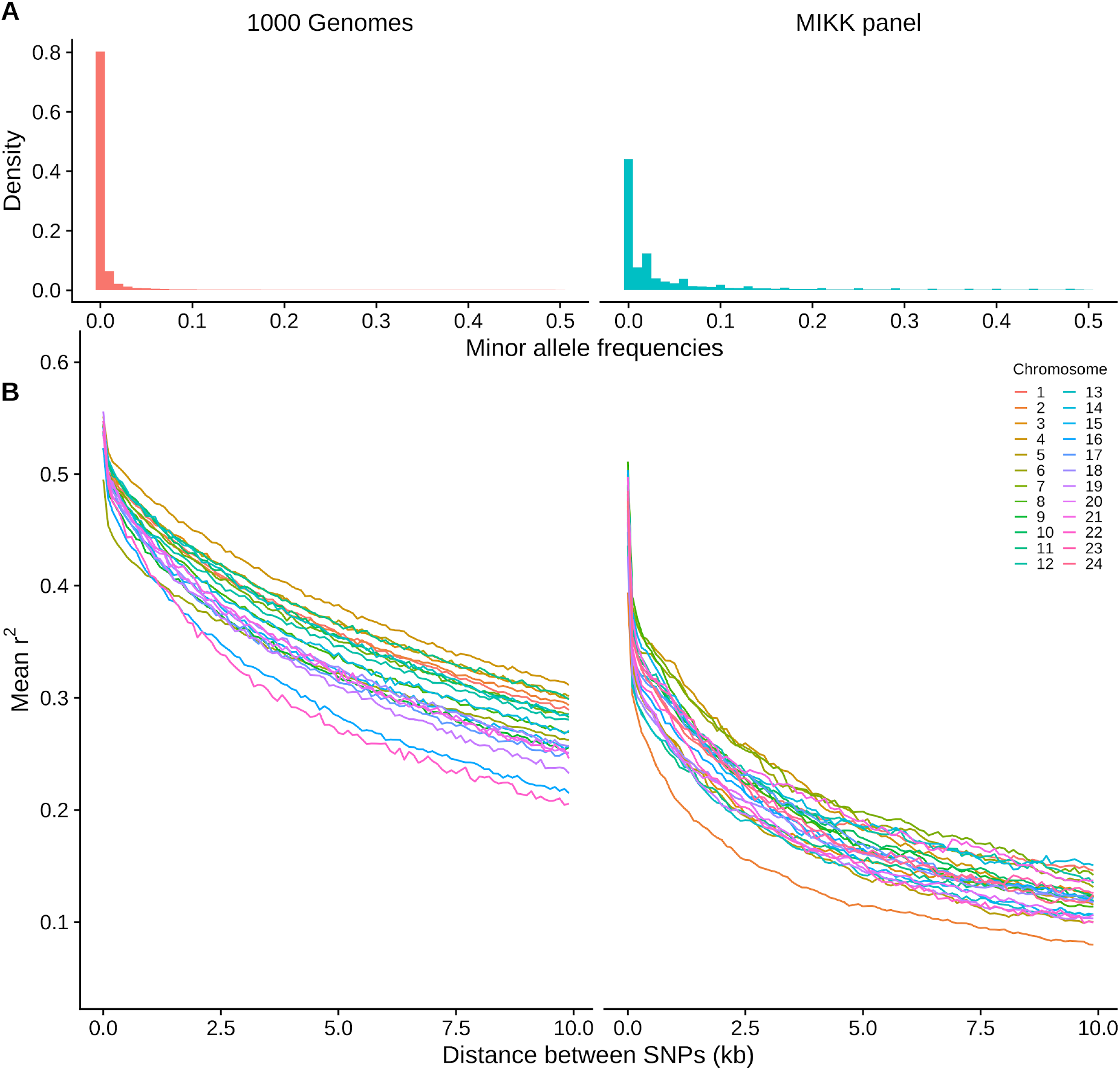
Minor allele frequency distributions and LD decay for biallelic, non-missing SNPs in the 1000 Genomes Phase 3 variant calls (N = 2,504) (1KG), and the MIKK panel Illumina-based calls excluding one of each pair of sibling lines (N = 63), across all autosomes (1KG: chrs 1-22; MIKK: chrs 1-24). **A**: Density plot of allele frequencies in the 1KG and MIKK panel calls. **B**: LD decay for each autosome, calculated by taking the mean r^2^ of pairs of SNPs with MAF > 0.1 within non-overlapping 100-bp windows of distance from one another, up to a maximum of 10 kb. Inset: mean r^2^ within 100-bp windows, up to a maximum of 1 kb. LD decays faster on chromosome 2 for the MIKK panel due to its higher recombination rate, consistent with the genetic linkage map described in [45].

To review the MIKK panel’s LD structure, for each autosome in humans (chromosomes 1-22) and each chromosome in medaka (chromosomes 1-24), we calculated *r*^*2*^ between all pairs of biallelic SNPs with a minor allele frequency (MAF) greater than 0.10, within 10 kb of one another (**Methods**). We then grouped the paired SNPs by distance from one another into non-overlapping 100-bp windows, and calculated the mean *r*^*2*^ for each window in order to represent how LD decays with distance between loci. **Figure 8B** compares the LD decay between the 1KG and MIKK panel SNPs. Based on the 1KG calls under these parameters, LD decays in humans to a mean *r*^*2*^ of around 0.2-0.35 at a distance of 10 kb, whereas the MIKK panel reaches this level within 1 kb, with a mean *r*^*2*^ of 0.3-0.4 at a distance of ~100 bp. This implies that when a causal variant is present in at least two lines in the MIKK panel, one may be able to map causal variants at a higher resolution than in humans. We note that LD decays faster in chromosome 2 of the MIKK panel relative to the other chromosomes. This suggests that it has a much higher recombination rate, which is consistent with the linkage map described in [45] showing a higher genetic distance per Mb for this chromosome. This higher recombination rate in chromosome 2 may in turn be caused by its relatively high proportion of repeat content (**Supplementary Figure 1**).

### Copy Number Variation in the MIKK panel

To call copy number variation (CNV) across all MIKK panel lines, we used high-depth Illumina sequencing data to generate relative copy number estimates using an internal dynamic reference made up of at least 20 MIKK panel samples (**Methods**). We then called CNVs across the MIKK panel jointly, using a Hidden Markov Model (HMM) specifically designed to call CNVs from short-read data (**Methods**). Overall, we detected a total of 106,861 CNVs genome-wide across the MIKK panel compared to the *HdrR* reference, consisting of 59,284 losses and 47,577 gains, the majority of which were observed at low frequency across the panel (**Supplementary Figure 2**). Similar to the LoF analysis, we used the pLI score from the Exome Aggregation Consortium (ExAC) database [36] to derive a logarithm of odds (LOD) score for each CNV. Here we calculated pLI LOD scores for every called CNV across the MIKK panel by mapping medaka genes to human orthologues (**Methods**). Although only 11,411 medaka genes (48% of coding genes [46]) mapped successfully to a human ortholog with a pLI score available, we were still able to get a good representation of likely dosage intolerance across the majority of the medaka genome (**Supplementary File 2**).

The majority of common CNVs do not fall in regions of high dosage intolerance for human orthologous genes across the medaka genome (**Figure 9A**), and as expected higher frequency CNVs tend to have lower pLI LOD scores (**Figure 9C**). CNV events with very large pLI LOD scores (> 20) are observed exclusively as either 1 or 3 copy events (**Figure 9B**), with those with more extreme copy numbers (zero or greater than 3 copies) showing a clear decrease in the number of positive pLI LOD scores (**Figure 9B**). Interestingly, there is little difference in the pLI LOD score distributions between losses compared to gains, with losses showing a marginal increase in the overall numbers of high pLI LOD scores. The vast majority of CNVs are low frequency across the MIKK panel, and higher frequency events are less likely to be observed in predicted dosage-sensitive genes, suggesting that there may have been a degree of purging of highly deleterious loss of function CNVs during the inbreeding process.

**Figure 9:**
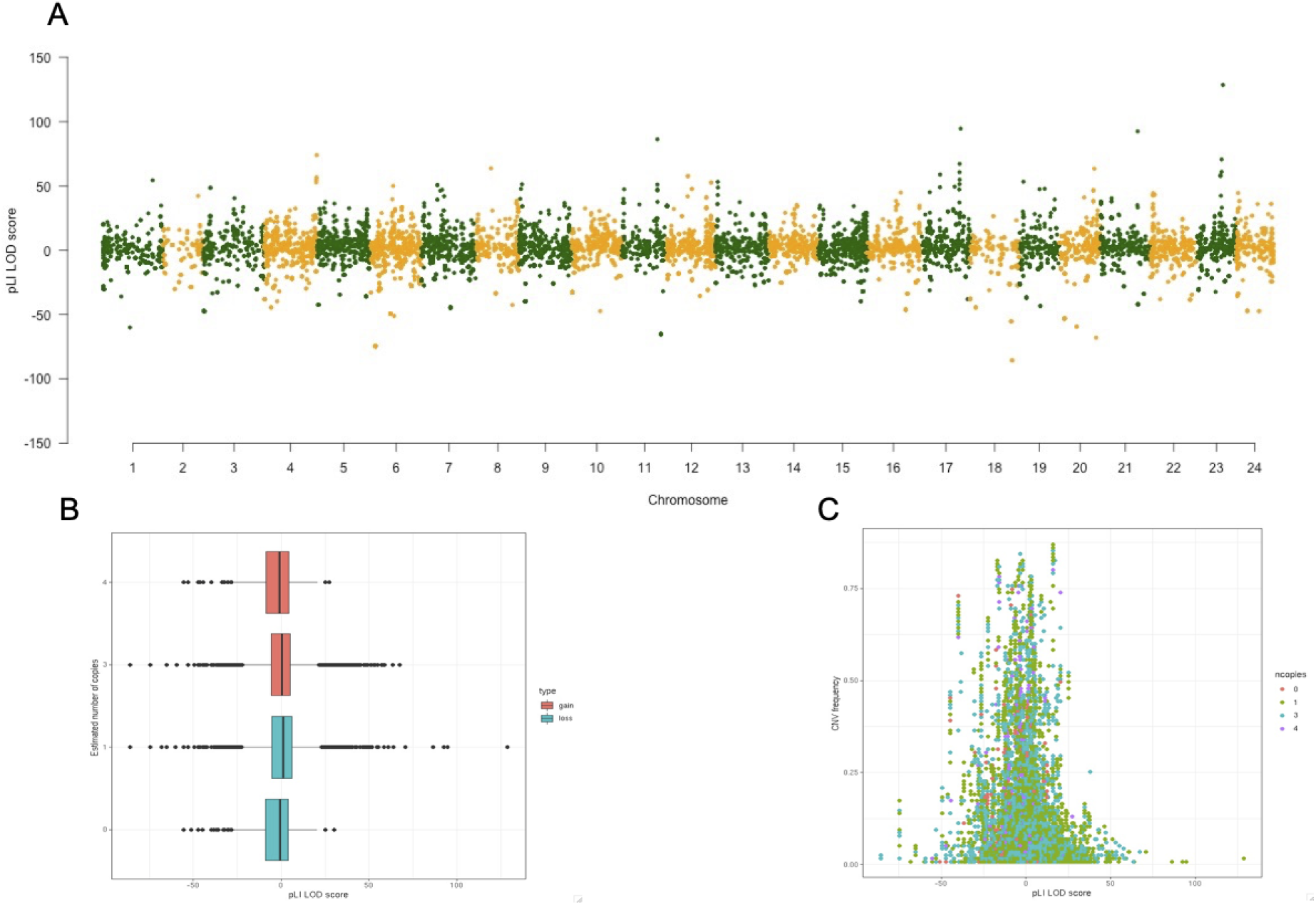
pLI LOD score for CNVs across the medaka genome. **A**: Genome-wide view of pLI LOD score. **B**: Distribution of pLI LOD score by the predicted number of copies for all CNVs. **C**: pLI LOD scores against CNV frequency coloured by the predicted number of copies for each CNV.

### Expression QTL analysis

To investigate the genetic diversity of the panel at the transcriptome level we performed a proof of concept expression-based quantitative trait analysis (eQTL) using liver tissue from 50 female MIKK panel lines (column G, **Supplementary Table 1**). We found 14,835 significant eQTL SNPs (*1% FDR*) in 3,795 transcripts corresponding to 3,543 genes (**Figure 10A, Supplementary File 3** and **Supplementary Table 6**) and observed a highly consistent pattern of expression between all samples. Interestingly, most of the sibling lines included in the RNA analysis clustered closely together, suggesting that broad-scale expression differences, as a direct consequence of the genomic sequence in medaka, are heritable across multiple generations (**Supplementary Figure 3**), and suggestive of broader trans-effects in expression. To further explore the genotypic pattern associated with expression profiles, we looked in detail at a stretch of 3 SNPs associated with an eQTL in an isoform of *cyp17a2* ([ENSORLT00000002786.2](https://www.ensembl.org/Oryzias_latipes/Transcript/Summary?db=core;g=ENSORLG00000002242;r=18:551864-558794;t=ENSORLT00000002786)) – a gene likely involved in progesterone metabolism (**Figure 10B,C**). For all 3 SNPs there is a strong correlation between the level of transcript expression and the genotype, with T genotype being associated with a much higher median expression, while A genotype has little to no expression.

**Figure 10:**
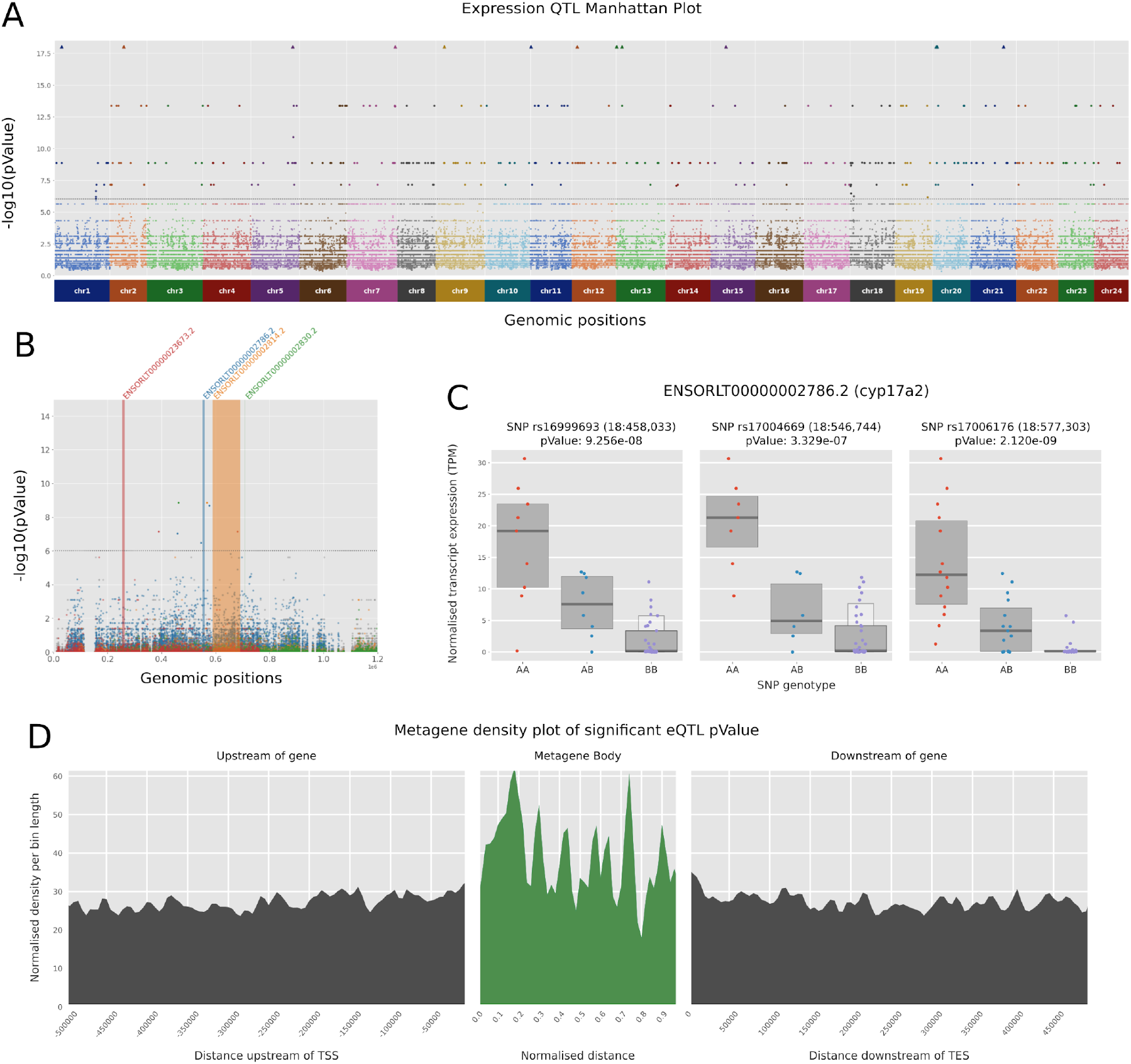
eQTL analysis results: **A**. Manhattan plot reporting only the 5 best p-values per transcript. For visual comfort, the p-value threshold (dashed line) was set to 1e-6. **B**. Zoomed-in slice of the Manhattan plot, showing all p-values in the genomic window chr8:0-1200000. Transcripts associated with at least 1 p-value lower than 10e-6 are highlighted in colour, with all SNPs associated shown in the same colour. The width of the coloured bands corresponds to the full length of the transcript. **C**. Normalised TPM expression of MIKK lines for 3 significant eQTL SNPs associated with transcript ENSORLT00000002786.2 (cyp17a2 gene). Lines are sorted according to their genotypes (AA, AB or BB). Each point represents one individual line, overlaid with letter-value plots boxes showing the quantile distribution [50]. **D**. Metagene density plot with upstream and downstream 500k regions showing the density of significant eQTL hits according to their positions relative to a prototypic gene. Values are aggregated in 100 bins of 5000 bases for the adjacent regions and 50 bins dynamically computed for p-values falling inside gene bodies. Densities are weighted by bin length to allow comparisons across the different regions.

Finally, we generated a metagene plot showing an aggregated distribution of the location of all significant eQTLs, showing a slight enrichment immediately downstream and upstream of genes and an enrichment peak inside gene bodies shortly after the transcription start site (TSS) (**Figure 10D**). This is consistent with variation in and around the TSS being likely to impact gene expression. Although noisy due to the small number of associations from this limited sample size, the pattern is similar to the previously reported eQTL metagene plot in vertebrates [47] (**Figure 10B)**. Altogether, we showed that despite the small number of samples, we are able to identify transcripts associated with eQTLs in the MIKK panel with similar properties to human and other vertebrate eQTL studies [48,49].

## Discussion

We have established a panel of 80 near-isogenic inbred lines of medaka from a carefully-selected wild population without prior genetic domestication of founder animals. In addition to the number of inbred lines, the MIKK panel is unique among vertebrate inbred panels as it has been derived from a wild population. It is well known that the domestication of wild animals often has severe consequences for the genetic diversity of a population [21,51]. In the MIKK panel, genetic adaptation to husbandry conditions prior to inbreeding did not occur. This design allowed us to preserve much of the naturally-occurring genetic variation in its wild founding population. We speculate that the tolerance to inbreeding from the wild is in part due to an ability of wild medaka to adapt to facility husbandry conditions. Consistent with this, medaka has a long history as a pet animal in Japan, demonstrating the natural propensity of this species to tolerate and reproduce in captivity [22]. The MIKK panel is therefore a unique tool to examine phenotype-genotype associations in a natural vertebrate population under precisely-controlled laboratory conditions. Our estimates of fecundity levels indicate that egg production is sufficient to meet the demands for the phenotyping of complex traits that often show quantitative variation, and therefore require the measurements of large numbers of individuals. In the future, carefully controlled quantification of fecundity levels will address whether the observed variation between lines has a genetic component.

We confirmed that the panel has captured phenotypic variability by exploring two phenotypes. First, for a morphological trait by image analysis, we observed a broad sense heritability (*H*^2^) measure of 0.68 for eye size as an extractable trait. Second, we performed RNA-Seq on female liver tissues. We detected 622 significant cis expression quantitative trait loci (cis-eQTLs) in 541 protein-coding genes, providing a proof of concept as the basis for future extended analyses. The location of cis-eQTLs with respect to gene structure shows the expected enrichment towards transcriptional start sites, although less pronounced than in previous studies involving larger sample sizes [48,52]. In summary, this indicates that genetically-determined phenotypic variation is well represented in the MIKK panel.

High-depth Illumina sequencing revealed a high degree of homozygosity across all panel genomes. We see a striking reduction in genome-wide heterozygosity with isogenic levels similar to that of classical inbred strains that have been inbred for more than 70 generations. The distribution of homozygosity across the genome shows no bias towards any particular genomic region, with the exception of chromosome 1. As expected, we detected the expected drop in homozygosity around the sex-determining locus. Interestingly, 3 male fish showed no heterozygosity across this region in the X chromosome. This is consistent with the known XX males observed in medaka fish [53], and is also consistent with the recent evolution of genetic sex determination in *Oryzias latipes.* We therefore speculate that phenotypic males of XX genomic configuration were sequenced in these three cases. These lines continue to produce balanced ratios of males and females at the time of writing (generation F18) and will be interesting for studying aspects of the medaka sex determination system in the future.

The MIKK panel originates from a single population of wild medaka caught in Kiyosu, Japan. When comparing the genome-wide allele frequency distribution between the inbred lines and wild Kiyosu medaka, we observed similar allele frequencies and no large regions with distortions. This suggests that the genetic variation in the MIKK panel is a good representation of wild genetic variation, making the panel an excellent resource for genome-wide association studies (GWAS) using natural standing variation. To assess the resolution for genetic association mapping we calculated the linkage disequilibrium (LD) across MIKK panel lines. We compared the LD decay to that observed in human cohorts (1000 Genomes Project Phase 3), and observed a marked increase in LD decay, indicating that when a causal variant is present in more than one MIKK panel line, the mapping resolution may be higher than in humans. We note the faster decay of LD in chromosome 2, which is consistent with the known high recombination rate seen across this chromosome relative to its physical distance (3.36 cM/Mb in chromosome 2 vs 1.02-2.76 cM/Mb for the other chromosomes) [45], possibly attributable to that chromosome’s high repeat content (**Supplementary Figure 1**). There are potential analogies with the high recombination frequency present across chromosome 19 in humans [54].

One interesting aspect of the panel design is the fact that some lines have a ‘sibling’ line (originating from the same founder F1 cross). This special genetic structure across the panel that can be exploited for specific research applications, for example to explore the aggregate impact of epistatic (GxG) interactions without locus-level resolution. This genetic structure also acted as a definitive control for us during this analysis. When we look at measures of relatedness across the panel, the IBD analysis showed that ‘sibling’ lines cluster closely together, indicating that they are indeed more related to each than other panel lines. This finding not only shows that the panel displays the favourable genetic structure that was designed from the outset, but also that all the crossing procedures during the inbreeding process were robust.

To place the MIKK panel in context of medaka populations genetics around Japan, we performed ABBA-BABA analysis to compare the level of admixture between the MIKK panel and southern Japanese (*iCab*), northern Japanese (*HNI*), and Korean (*HSOK*) medaka strains. As expected, the MIKK panel shows the most admixture with the southern strain *iCab*, as Kiyosu is in southern Japan. However, there is a slight increase in admixture proportion for *HSOK* compared to *HNI*. Although the proportional difference is small, this supports the general finding that northern and southern Japanese medaka strains show low levels of interbreeding that may be a result of geographical isolation or genome divergence [55].

The majority of MIKK panel genomes are highly isogenic and as expected we observe low numbers of predicted loss-of-function (LoF) variants (<0.3% of all variants). Small homozygous LoF variants (SNPs and INDELs) show a slight enrichment in regions of residual heterozygosity in MIKK panel lines. This may indicate a purging of LoF variation in essential dosage-sensitive genes, although the observed enrichment is slight, which is likely due to the low change of LoF variation occurring in essential genes across the 9 generations of inbreeding, given the very low rate of spontaneous mutations. For CNVs, we see low numbers of large events across the MIKK genomes. Large CNVs are likely to impact on several genes and can often result in LoF for multiple genes at once, in particular for losses compared to gains [56]. An assessment into the CNV landscape across the MIKK panel genomes in relation to dosage sensitivity revealed that the majority of events occur outside of dosage sensitive regions, and that potentially impactful CNVs encompassing several dosage sensitive genes are seen at very low frequency. There are, however, many rare genetic variants (both SNVs and CNVs) that occur in genes that are predicted to have a functional impact or an intolerance to dosage. This is, of course, not unexpected but it does highlight one of the powerful applications of genetic screening using the MIKK panel, where specific F2-cross designs can be used to balance the distribution of rare impactful variants across an F2 population, allowing a functional analysis of rare variants during trait association analyses. Exploration of the phenotypes of these cases, or mapping of potential modifier mutations that might compensate for such loss of function variation, will be a valuable line of research for both the medaka and human disease communities.

Here we provide a detailed analysis of some of the sources of variation present across the MIKK panel lines, and put in place important resources to allow the MIKK panel to be fully exploited during further research studies. Further sequencing of the MIKK panel lines, using both long and short read technologies is planned in the future as well as extensive molecular, organ function and organismal phenotyping across the panel. In a separate analysis [57], we looked at building line-specific genome reference datasets from a subset of the MIKK panel, and show that by using non-linear reference alignment approaches (graph genomes), we are able to build a more complete representation of genetic variation in the MIKK panel, uncovering additional variation that is masked when using standard reference-alignment approaches. Further detailed characterization of the MIKK panel in terms of genetic variation as well as molecular, organ and organismal phenotyping will provide a wealth of information allowing us to leverage the isogenic properties of the panel to investigate phenotype-to-genotype interactions (including gene-to-environment effects, or GxE) at high genetic resolution. We invite the medaka, telost and broader vertebrate genetics community to make use of the resources presented here, and to contact the authors to explore the hosting of experiments across the MIKK panel in the style of a “research hotel”.

## Acknowledgements

The authors acknowledge Thomas Seitz, and Irina Oks and Marzena Majewski for supporting the fish husbandry, and Alicia Günthel, Rachel Müller and Beate Wittbrodt for laboratory assistance. F.L. dedicates this paper to Sabine and Rolf Loosli-Walther. The work was funded by the Helmholtz funding programme BIFTM to F. Loosli, N. Wolf, N. Kusminski, C. Herder and N. Aadepu. E. Birney, T. Fitzgerald, A. Leger, C. Barton, J. Monahan and I. Brettell were funded by the EMBL European Bioinformatics Institute (EMBL-EBI). This work was supported by Heidelberg University Core Funding to T. Tavhelidse, T. Thumberger and J. Wittbrodt. This project has received funding from the European Research Council (ERC) under the European Union’s Horizon 2020 research and innovation programme (grant agreement No 810172), from the NIH UH-3338-03 (JW), the German Ministry for Research (BMBF: HIGH-life 05K19VH1, Code-Vita 05K16VH1, JW) and the German Center for Heart Diseases DZHK (JW, JG).

## Author contribution

Conception of the project: E.B., T.F., F.L., K.N., J.W.; Project management and supervision: E.B., T.F., F.L., J.W.; Sampling of wild fish: K.N.; Inbreeding: N.K., F.L., N.W.; Sample preparation: N.A., C.B., J.G., C.H., E.H., O.T.H., C.L., K.L., F.L., R.S., E.T., T.Ta., T.Th., P.W., B.W., J.W.; Image acquisition: N.A., C.H., N.Kh.; Data analysis: C.B., E.B., I.B., T.F., A.L., F.L., J.M., J.W.; Manuscript writing: E.B., I.B., T.F., A.L., F.L., J.W.

## Materials and Methods

### MIKK husbandry and inbreeding

Medaka (*Oryzias latipes*) fish were maintained at the medaka facility of the KIT Institute of Biological and Chemical Systems, Biological Information Processing (IBCS-BIP). All animal husbandry and experimental procedures were carried out in accordance with local and European Union animal welfare standards (Tierschutzgesetz §11, Abs. 1, Nr. 1, AZ35-9185.64/BH). The facility is under the supervision of the local representative of the animal welfare agency.

An unstructured polymorphic wild population of medaka was sampled in the irrigation canals of Kiyosu near Toyohashi as described [31]. 141 random single pair mating crosses of wild Kiyosu fish were set up at NIBB, Okazaki, and their F1 offspring shipped to the medaka facility of KIT. Of these 141 crosses, the surviving 115 founder F1 families were raised to adulthood and used for 9 generations of single full-sibling-pair inbreeding to establish the panel of near-isogenic lines now known as the Medaka Inbred Kiyosu-Karlsruhe (MIKK) panel.

Full-sibling-pair inbreeding crosses (Hyodo-Takuchi 1980) were set up as follows: approximately 1 month after the fishes (siblings) of a line started productive mating, random single brother-sister crosses were set up in 2-litre mating cages with water recirculation. In crosses resulting in only unfertilized eggs, the male was exchanged with a male of the same strain (i.e. brother). Eggs were collected from females after 4 days of successful mating. Unproductive crosses were aborted after 2 weeks and a new single brother-sister (full-sibling-pair inbreeding) cross of the same line was set up. Eggs were collected from a given mating cross until at least 50 hatchlings were obtained that survived for 2 weeks.

MIKK panel fish at the medaka facility of KIT were kept as described in [58], in Müller & Pfleger systems (40 6-liter tanks per recirculation system; Müller & Pfleger, Rockenhausen; Germany) with the following modifications: eggs were raised at 22°C. Under the constant summer conditions (14 hours light/10 hours dark: 14:10LD) medaka mate every day, mostly at the onset of the light period [32]. A *flavobacterium columnare* infection that occurred while inbreeding generations 4-6 was treated with Baytril (Baytril Enrofloxacine 10% injection solution, Bayer; 2ml/100l system water for 7 days followed by a 50% system water exchange). Thereafter the inbreeding lines were kept at 23°C to reduce the microbial load in the system water.

### Imaging of adults and image analysis

to 9-month-old adults were sacrificed by hypothermic shock. Lateral and dorsal images were taken with a Canon EOS 250D camera and a Sigma 18-250mm lens. Using a subset of the MIKK panel images, a number of machine learning models were trained to detect and measure phenotypes from the images of the entire panel of fish. The code for this was developed in Python using an existing implementation of semantic image segmentation in the Keras (https://github.com/divamgupta/image-segmentation-keras) machine learning framework with Tensorflow [59] used as the backend.

The network architecture used in these models was a semantic segmentation network known as SegNet. This is a deep fully convolutional neural network architecture for semantic pixel-wise segmentation. The SegNet architecture is based on a modification of the VGG16 network proposed which makes the network more lightweight and quicker to train. This architecture was chosen primarily as it has been shown to improve boundary detection and reduce the number of parameters that need optimisation, allowing for easier end-to-end network training. Additionally, this architecture has been shown to achieve high training accuracy when training with small datasets.

A subset of the panel images were segmented using the Labelbox platform [60] to generate the training data to train models to segment the fish body, eye, tail and anal fin within the images. During training the training dataset was augmented using the Python library imgaug [61]. The following affine transformation where randomly applied to the images during training:

- scale images to 80-120% of their size,
- translate images by −20 to +20 relative to height/width (per axis),
- rotate images by −45 to +45 degrees,
- shear images by −16 to +16 degrees.

After training the models they were used to predict the chosen features in the remaining unseen images and the following parameters were extracted from these segmentations.

- nose to tail length in pixels,
- maximum width of fish in pixels,
- fish area in pixels,
- eye area in pixels,
- length of abdominal region in pixels,
- length of caudal region in pixels,
- ratio between lengths of the abdominal and caudal region,
- eye diameter in pixels,
- distance from the top of the eye to the top of the head,
- distance from the bottom of the eye to the bottom of the head.

### Tissue dissection

For whole genome sequencing, medaka organs were dissected from 6-month-old male adults. Fish were sacrificed by hypothermic shock. The brain was dissected and shock frozen in liquid nitrogen. For RNAseq analysis 12 month old adults that were kept at either 14 light:10 dark (summer condition) or 10 light:14 dark (winter condition) light cycles respectively were sacrificed by hypothermic shock and the organs after dissection were shock frozen in liquid nitrogen.

### DNA and RNA extraction

#### DNA extraction and whole Genome Sequencing of the MIKK panel

DNA was extracted from medaka brains in 2ml Eppendorf tubes using the Qiasymphony DSP DNA Mini kit (Cat. No. 937236). Tissue_HC_200_v7 Protocol, 100ul elution volume. Pre-treatment samples were homogenised with 5mm stainless steel beads in 220ul buffer ATL at 30Hz for 20 seconds on the TissueLyser. 20μl proteinase K was added, and samples were incubated for 1h at 56C, 900rpm on the Qiacube (an alternative heater-shaker platform would be an Eppendorf Thermomixer). RNase treatment, 2min incubation at RT. Following DNA extraction from whole brain samples, each line of the MIKK panel was whole genome sequenced at greater than 30x coverage using Illumina X10 instrumentation at the Wellcome Trust Sanger Institute. Library preparation was performed following the standard PCR-free Illumina protocol [62]. 1ug was picked from DNA extraction plates within the Sanger sample logistics facility and passed into the Sanger sequencing pipeline from library preparation. Following successful preparation of sequencing libraries, the samples were QCed using Qubit and samples passing the facility quality control threshold were multiplexed sequenced in paired end mode with 5 samples per Illumina X10 flow cell.

#### RNA extraction and Illumina RNA-sequencing

RNA extraction from 52 liver samples was performed on a Qiagen automated extraction platform using QIAsymphony RNA Kits, where polyA RNA was extracted from liver samples for paired-end RNA-Seq analysis. Samples were prepared for Illumina RNA-Sequencing using the NEBNext Ultra II Directional RNA Library Prep Kit for Illumina and sequenced on a Hiseq 4000 sequencing platform following the manufacturer’s instructions. We obtained data passing the facility quality control threshold for 50 samples out of the 52 initially processed.

#### Bioinformatic methods and data

Raw sequencing data can be retrieved from ENA linked to the following project ID:

- Illumina DNA sequencing data: PRJEB17699
- Illumina RNA sequencing data: PRJEB43091

All the scripts and metadata used for this study are extensively described in the associated github repository available at https://github.com/birneylab/MIKK_genome_main_paper

#### DNA Sequence Alignment and SNP calling

After sequencing, the MIKK panel genomic data was transferred securely from Sanger compute storage to EBI where alignment and variant calling was performed. All whole genome sequence (WGS) datasets were aligned against the latest medaka *HdrR* reference genome from ENSEMBL (release 94) using the Burrow-Wheeler Aligner for short-read alignment (BWA) [63]. For variant-calling we used the best practices approach for calling SNPs and INDELs using the GATK software [64]. Briefly, aligned BAM files were processed using the Picard software [65] to make duplicate reads and correct read group tags prior to the creation of gVCF files using the HaplotypeCaller command from GATK. Finally, gVCFs were combined and genotyped using GenotypeGVCFs from GATK. The result of this processing was a single multisample VCF file containing SNP and INDEL calls across all lines of the MIKK panel (MIKK Illumina callset).

#### RNA sequencing data processing and eQTL analysis

We developed a Snakemake pipeline [66] called NanoSnake [67] to run the entire analysis, including read trimming, mapping, quality control and transcript abundance estimate. For this study we ran NanoSnake v0.0.3.1 RNA_Illumina workflow (https://github.com/a-slide/pycoSnake/tree/0.0.3.1). All the tools and environment are version-controlled in individual conda environments. Briefly, reference genome, transcriptome and annotations were obtained from ensembl Release 98 (Japanese medaka *HdrR* ASM223467v1, https://www.ensembl.org/Oryzias_latipes/Info/Index). For each of the 50 datasets (column G, **Supplementary Table 1**), Illumina pair-end reads were cleaned-up using Fastp v0.20.0 [68], aligned to the reference genome using STAR v2.7.3a [69] and obtained estimated counts per gene. In parallel, we also performed a transcriptome pseudo-alignment transcript level quantification using Salmon v1.1.0 [70]. The configuration file and run script used for NanoSnake are available at https://github.com/birneylab/MIKK_genome_main_paper/tree/master/eQTL/code/nanosnake.

We estimated transcript abundances with Salmon [70] and ran the eQTL analysis with Limix [71], looking for Cis-eQTLs within 250 kb of each transcript and using the genetic relatedness matrix between lines (**Supplementary Figure 4**) as a covariate.

For the eQTL analysis, we used the transcript abundance estimates obtained from Salmon and applied a pre-filtering step to retain transcripts with a minimal count of 5 reads and a minimal TPM (transcript per million) of 1 in at least half of the samples (30/60). For valid transcripts (13413 out of 36777) we then normalised the TPM values using a quantile Gaussian normalisation. SNP genotypes were cleaned-up and preprocessed with PLINK [72]. We performed a light LD pruning to filter out highly correlated variants, retaining 45.7% of all variants. Using limix v3.0.4 [71] we first computed a genetic relatedness (kinship) matrix. Then, for each valid transcript we ran a LMM univariate association test between the observed phenotypes (normalized TPM) and the genotypes of every SNPs within 500 kb upstream and downstream to the transcript, using the kinship matrix as a covariate (limix.qtl.scan). Finally, we used python scripting to extract significant hits and generate visualizations, including Manhattan and metagene plots. The entire analysis notebook and raw data are available at https://github.com/birneylab/MIKK_genome_main_paper/tree/master/eQTL.

#### Definition of Homozygous Blocks across the medaka genome

To assess the level of homozygosity in the MIK panel we applied a number of different approaches. Firstly, we used the PLINK software [72] to calculate the inbreeding coefficient (*F*) from the MIKK panel genotype calls. Additionally we used PLINK to call runs of homozygosity across each of the MIKK panel lines across the medaka genome, as well as calculating identity by descent (IBD) of all panel lines. To estimate an accurate level of homozygosity and create a homozygous block call set across the entire medaka genome we developed a Hidden Markov Model (HMM) to call the allelic status across all MIKK panel lines using 10 kilobase (kb) resolution across the medaka genome. Briefly, the number of heterozygous SNP genotype calls were counted across 10-kb non-overlapping windows in the medaka genome for each MIKK panel line separately. These genotype counts were used to train a 2-state HMM with a Poisson distribution using the expectation maximisation (EM) algorithm, followed by the Viterbi (forward/backward) algorithm to define the most likely state space path. Following this process, for each MIKK panel line, we have 10 kb regions of the medaka genome that were called as either heterozygous or homozygous.

#### Copy number detection (CNV) from Illumina sequence data

To call CNVs across the medaka genome in all MIKK panel lines, we used a method based on read depth and cross-sample normalisation to generate robust copy number estimates (log_2_ ratio values) prior to using a specifically developed HMM to call copy number variable regions across the genome.

First, read depth information was extracted from alignment BAM files at 2-kb non-overlapping windows across the medaka genomes. Reads that were not proper pairs or where either read mate was unmapped were excluded, as were PCR duplicates or reads containing secondary alignments. Next, to generate sample-specific dynamic-baseline references, an all-by-all correlation test was performed to group samples based on read depth similarity, prior to transforming read depth values to normalised log_2_ ratio values based on the median depth across its most highly correlated samples (*n*=20). Next, a three-state HMM was trained on all data in combination prior to detecting variable segments using the Viterbi algorithm [73].

#### Population genetic analysis of the MIKK panel

First, we wanted to test the genotypes contained in the MIKK panel lines against the distribution of alleles from the original wild population of Kiyosu medaka. To do this we calculated the fixation index (*F*_*ST*_) using genotype information from 8 wild Kiyosu lines, comparing the allele frequencies against those in the MIKK panel.

#### Introgression with Northern and Korean medaka strains

In light of previous findings [31], we sought to compare the extent to which the MIKK panel showed evidence of introgression with established inbred medaka strains originating from southern Japan (*iCab*), northern Japan (*HNI*), and Korea (*HSOK*). We used the 50-fish multiple alignment from Ensembl release 102 [74] to obtain the aligned genome sequences for *HdrR*, *HNI* and *HSOK*, as well as the most recent common ancestor of all three strains. Using the phylogenetic tree provided with the dataset, and the *ape* R package version 5.4.1 [75], we determined the most recent common ancestor of those three strains. For each locus with a non-missing base for *HdrR*, we assigned the allele in that ancestral sequence as the ‘ancestral’ allele, and the alternative allele as the ‘derived’ allele, and then combined that dataset with the Illumina VCF containing SNPs called in the MIKK panel lines and *iCab* strain. We then calculated 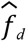 in sliding windows as described in [42], using the scripts provided by the first author on their GitHub page (https://github.com/simonhmartin/genomics_general). Data processing and analyses were carried out with R version 4.0.2 [76], and the Tidyverse suite of R packages version 1.3.0 [77]. Figure 7 was generated using the R packages *ape [75]* and *circlize [78]*.

#### MAF and LD decay in the MIKK panel

The MIKK Illumina callset contained SNP and INDEL calls for 79 of the 80 extant MIKK panel lines (column D, **Supplementary Table 1**). In order to avoid allele frequency biases introduced by the 16 pairs/triplets of “sibling lines”, we removed each pair’s arbitrarily-labelled second sibling line from the callset, leaving 63 MIKK panel lines for this analysis (column H, **Supplementary Table 1**) (‘MIKK non-sibling calls’). To assess how accurately one may be able to map genetic variants using the MIKK panel relative to a human dataset, we compared the MIKK panel’s minor allele frequency (MAF) distribution and LD structure against that of the 2,504 humans in the 1KG Phase 3 release [44]. To prepare the ‘1KG calls’, we first downloaded the VCFs for each autosome from the project’s FTP site (ftp://ftp.1000genomes.ebi.ac.uk/vol1/ftp/release/20130502/), then merged them into a single VCF using GATK [64].

We used PLINK [79,80] to calculate the minor allele frequencies for all non-missing, biallelic SNPs in both the MIKK non-sibling and IKG calls (*N*_SNPs_ = 16,395,558 and 81,042,381 respectively), and then used R [81] and the tidyverse suite of R packages [77] to produce Figure 8A. To visualise the MIKK panel’s LD structure, we first used PLINK to filter the MIKK non-sibling callset for non-missing, biallelic SNPs with MAF ≥ 0.10, leaving 2,968,786 SNPs. We then used PLINK to take a random sample of 3000 SNPs per chromosome and recode them for analysis in Haploview [82]. We used Haploview to generate LD plots covering the length of each of the 24 chromosomes in the default colour scheme, showing *r*^*2*^ values for each pair of SNPs within 1 Mb of each other (**Supplementary File 4**). To determine the rate of LD decay in the MIKK panel and compare it to that in the 1KG sample, for both the MIKK non-sibling calls and the 1KG calls, we used PLINK to compute *r*^*2*^ on each autosome for all pairs of non-missing, biallelic SNPs with MAF > 0.10 within 10 kb of one another (for 1KG and the MIKK panel respectively: ~5.5M and ~3M SNPs, with a total number of pairwise *r*^*2*^ observations of 204,152,922 and 146,785,673). We then used R and the tidyverse suite of R packages to group the *r*^*2*^ observations for each pair of SNPs based on their distance from one another into non-overlapping bins of 100-bp in length, then calculated the mean *r*^*2*^ in each of those bins and generated the plots in Figure 8B using the mean *r*^*2*^ and left boundary of each bin.

#### Prediction and annotation of repetitive and transposable elements

The *RepeatModeler* pipeline (v2.0.0) [83] for the automated *de novo* identification of repetitive and transposable elements was run on all chromosomes in the *HdrR* genome assembly [84]. RepeatModeler was run with its default parameters and the additional long terminal repeat (LTR) structural discovery sub-pipeline that includes the *LTRharvest [85]* and *LTR_retriever [86]* tools.

The RepeatModeler library of repeats was filtered to remove non-TE protein coding sequences by using a protein BLAST (Altschul *et al.*, 1990) to align (*E-value* ≤ 1e-5) the *Oryzias latipes* proteome (Ensembl v99) and *pfam* peptide database (v32) against the RepeatMasker peptide library. Finally, a nucleotide BLAST was used to remove any RepeatModeler repeats that aligned (*E-value* ≤ 1e-10) against the corresponding transcripts.

RepeatMasker (v4.1.0) [87] was used to align the chromosomes in the *HdrR* assembly against the filtered RepeatModeler library of consensus repeats and the existing RepeatMasker repeat families.

Additionally, *Exonerate* (Slater and Birney, 2005) was used to align the two subtypes of the *Teratorn* mobile element found in the *Oryzias latipes* genome against the *HdrR* reference. (The *Teratorn* element being the result of a fusion between a *piggyBac* DNA transposon and a member of the *Alloherpesviridae* family [88]).

## References

1. Fisher RA. XV.—The Correlation between Relatives on the Supposition of Mendelian Inheritance. Earth Environ Sci Trans R Soc Edinb. 1919;52: 399–433.

2. Toyama K. On some Mendelian characters (in Japanese). Rep Jap Breed Soc. 1916;1: 1–9.

3. Alonso-Blanco C, Aarts MGM, Bentsink L, Keurentjes JJB, Reymond M, Vreugdenhil D, et al. What has natural variation taught us about plant development, physiology, and adaptation? Plant Cell. 2009;21: 1877–1896.

4. Mackay TFC, Huang W. Charting the genotype-phenotype map: lessons from the Drosophila melanogaster Genetic Reference Panel. Wiley Interdiscip Rev Dev Biol. 2018;7. doi:10.1002/wdev.289

5. Mackay TFC, Richards S, Stone EA, Barbadilla A, Ayroles JF, Zhu D, et al. The Drosophila melanogaster Genetic Reference Panel. Nature. 2012;482: 173–178.

6. Wellcome Trust Case Control Consortium. Genome-wide association study of 14,000 cases of seven common diseases and 3,000 shared controls. Nature. 2007;447: 661–678.

7. Astle WJ, Elding H, Jiang T, Allen D, Ruklisa D, Mann AL, et al. The Allelic Landscape of Human Blood Cell Trait Variation and Links to Common Complex Disease. Cell. 2016;167: 1415–1429.e19.

8. Ganna A, Verweij KJH, Nivard MG, Maier R, Wedow R, Busch AS, et al. Large-scale GWAS reveals insights into the genetic architecture of same-sex sexual behavior. Science. 2019;365. doi:10.1126/science.aat7693

9. Mackay TFC, Stone EA, Ayroles JF. The genetics of quantitative traits: challenges and prospects. Nat Rev Genet. 2009;10: 565–577.

10. Bergelson J, Roux F. Towards identifying genes underlying ecologically relevant traits in Arabidopsis thaliana. Nat Rev Genet. 2010;11: 867–879.

11. Visscher PM, Naomi WR, Zhang Q, Sklar P, McCarthy MI, Brown MA, et al. 10 Years of GWAS Discovery: Biology, Function, and Translation. Am J Hum Genet. 2017;101: 5–22.

12. Tam V, Patel N, Turcotte M, Bossé Y, Paré G, Meyre D. Benefits and limitations of genome-wide association studies. Nat Rev Genet. 2019;20: 467–484.

13. Korte A, Farlow A. The advantages and limitations of trait analysis with GWAS: a review. Plant Methods. 2013;9: 1–9.

14. The Complex Trait Consortium. The Collaborative Cross, a community resource for the genetic analysis of complex traits. Nat Genet. 2004;36: 1133–1137.

15. A platform for experimental precision medicine: The extended BXD mouse family. Cell Systems. 2021;12: 235–247.e9.

16. Threadgill DW, Miller DR, Churchill GA, de Villena FP-M. The collaborative cross: a recombinant inbred mouse population for the systems genetic era. ILAR J. 2011;52: 24–31.

17. Rosenthal N, Brown S. The mouse ascending: perspectives for human-disease models. Nat Cell Biol. 2007;9: 993–999.

18. Schofield PN, Hoehndorf R, Gkoutos GV. Mouse genetic and phenotypic resources for human genetics. Human Mutation. 2012. pp. 826–836. doi:10.1002/humu.22077

19. Morse HC III. Origins of Inbred Mice. Elsevier; 2012.

20. Wade CM, Daly MJ. Genetic variation in laboratory mice. Nat Genet. 2005;37: 1175–1180.

21. Larson G, Burger J. A population genetics view of animal domestication. Trends Genet. 2013;29: 197–205.

22. Takeda H, Shimada A. The art of medaka genetics and genomics: what makes them so unique? Annu Rev Genet. 2010;44: 217–241.

23. Aida T. On the Inheritance of Color in a Fresh-Water Fish, APLOCHEILUS LATIPES Temmick and Schlegel, with Special Reference to Sex-Linked Inheritance. Genetics. 1921;6: 554–573.

24. Kirchmaier S, Naruse K, Wittbrodt J, Loosli F. The genomic and genetic toolbox of the teleost medaka (Oryzias latipes). Genetics. 2015;199: 905–918.

25. Gutierrez-Triana JA, Tavhelidse T, Thumberger T, Thomas I, Wittbrodt B, Kellner T, et al. Efficient single-copy HDR by 5’ modified long dsDNA donors. Elife. 2018;7. doi:10.7554/eLife.39468

26. Vilella AJ, Severin J, Ureta-Vidal A, Heng L, Durbin R, Birney E. EnsemblCompara GeneTrees: Complete, duplication-aware phylogenetic trees in vertebrates. Genome Res. 2009;19: 327–335.

27. Howe K, Clark MD, Torroja CF, Torrance J, Berthelot C, Muffato M, et al. The zebrafish reference genome sequence and its relationship to the human genome. Nature. 2013;496: 498–503.

28. Hyodo-Taguchi - Zool. Mag.(Tokyo) Y, 1980. Establishment of inbred strains of the teleost, Oryzias latipes. ci.nii.ac.jp. 1980. Available: http://ci.nii.ac.jp/naid/10005820467/

29. Murata K, Kinoshita M, Naruse K, Tanaka M, Kamei Y. Medaka: Biology, Management, and Experimental Protocols. John Wiley & Sons; 2019.

30. Hyodo-Taguchi Y. Inbred strains of the medaka, Oryzias latipes (Development of Medaka Biology in Japan-Part I). The fish biology journal Medaka. 1996;8: 11–14.

31. Spivakov M, Auer TO, Peravali R, Dunham I, Dolle D, Fujiyama A, et al. Genomic and phenotypic characterization of a wild medaka population: towards the establishment of an isogenic population genetic resource in fish. G3. 2014;4: 433–445.

32. Koger CS, Teh SJ, Hinton DE. Variations of light and temperature regimes and resulting effects on reproductive parameters in medaka (Oryzias latipes). Biol Reprod. 1999;61: 1287–1293.

33. Nanda I, Kondo M, Hornung U, Asakawa S, Winkler C, Shimizu A, et al. A duplicated copy of DMRT1 in the sex-determining region of the Y chromosome of the medaka, Oryzias latipes. Proc Natl Acad Sci U S A. 2002;99: 11778–11783.

34. Matsuda M, Nagahama Y, Shinomiya A, Sato T, Matsuda C, Kobayashi T, et al. DMY is a Y-specific DM-domain gene required for male development in the medaka fish. Nature. 2002;417: 559–563.

35. Otake H, Shinomiya A, Kawaguchi A, Hamaguchi S, Sakaizumi M. The medaka sex-determining gene DMY acquired a novel temporal expression pattern after duplication of DMRT1. Genesis. 2008;46: 719–723.

36. Lek M, Karczewski KJ, Minikel EV, Samocha KE, Banks E, Fennell T, et al. Analysis of protein-coding genetic variation in 60,706 humans. Nature. 2016;536: 285–291.

37. Fuller ZL, Berg JJ, Mostafavi H, Sella G, Przeworski M. Measuring intolerance to mutation in human genetics. Nat Genet. 2019;51: 772–776.

38. Nagylaki T. Fixation indices in subdivided populations. Genetics. 1998;148: 1325–1332.

39. Holsinger KE, Weir BS. Genetics in geographically structured populations: defining, estimating and interpreting F(ST). Nat Rev Genet. 2009;10: 639–650.

40. Green RE, Krause J, Briggs AW, Maricic T, Stenzel U, Kircher M, et al. A draft sequence of the Neandertal genome. Science. 2010;328: 710–722.

41. Durand EY, Patterson N, Reich D, Slatkin M. Testing for ancient admixture between closely related populations. Mol Biol Evol. 2011;28: 2239–2252.

42. Martin SH, Davey JW, Jiggins CD. Evaluating the Use of ABBA–BABA Statistics to Locate Introgressed Loci. Mol Biol Evol. 2014;32: 244–257.

43. Details on a Compara analysis. [cited 7 Oct 2020]. Available: http://apr2020.archive.ensembl.org/info/genome/compara/mlss.html?mlss=1828

44. 1000 Genomes Project Consortium, Auton A, Brooks LD, Durbin RM, Garrison EP, Kang HM, et al. A global reference for human genetic variation. Nature. 2015;526: 68–74.

45. Naruse K, Fukamachi S, Mitani H, Kondo M, Matsuoka T, Kondo S, et al. A detailed linkage map of medaka, Oryzias latipes: comparative genomics and genome evolution. Genetics. 2000;154: 1773–1784.

46. Howe KL, Achuthan P, Allen J, Allen J, Alvarez-Jarreta J, Amode MR, et al. Ensembl 2021. Nucleic Acids Res. 2021;49: D884–D891.

47. Veyrieras J-B, Kudaravalli S, Kim SY, Dermitzakis ET, Gilad Y, Stephens M, et al. High-resolution mapping of expression-QTLs yields insight into human gene regulation. PLoS Genet. 2008;4: e1000214.

48. Strunz T, Grassmann F, Gayán J, Nahkuri S, Souza-Costa D, Maugeais C, et al. A mega-analysis of expression quantitative trait loci (eQTL) provides insight into the regulatory architecture of gene expression variation in liver. Sci Rep. 2018;8: 5865.

49. Mason VC, Schaefer RJ, McCue ME, Leeb T, Gerber V. eQTL discovery and their association with severe equine asthma in European Warmblood horses. BMC Genomics. 2018;19: 581.

50. Hofmann H, Wickham H, Kafadar K. Letter-Value Plots: Boxplots for Large Data. Journal of Computational and Graphical Statistics. 2017. pp. 469–477. doi:10.1080/10618600.2017.1305277

51. Geiger M, Sánchez-Villagra MR, Lindholm AK. A longitudinal study of phenotypic changes in early domestication of house mice. R Soc Open Sci. 2018;5: 172099.

52. Ikeda D, Koyama H, Mizusawa N, Kan-No N, Tan E, Asakawa S, et al. Global gene expression analysis of the muscle tissues of medaka acclimated to low and high environmental temperatures. Comp Biochem Physiol Part D Genomics Proteomics. 2017;24: 19–28.

53. Naruse K, Tanaka M, Takeda H. Medaka: A Model for Organogenesis, Human Disease, and Evolution. Springer Science & Business Media; 2011.

54. Kong A, Gudbjartsson DF, Sainz J, Jonsdottir GM, Gudjonsson SA, Richardsson B, et al. A high-resolution recombination map of the human genome. Nat Genet. 2002;31: 241–247.

55. Katsumura T, Oda S, Hiroshi M, Oota H. Medaka population genome structure and demographic history described via genotyping-by-sequencing. doi:10.1101/233411

56. Rodriguez-Revenga L, Mila M, Rosenberg C, Lamb A, Lee C. Structural variation in the human genome: the impact of copy number variants on clinical diagnosis. Genet Med. 2007;9: 600–606.

57. Leger A, Brettell I, Monahan J, Barton C, Wolf N, Kusminski N, et al. Genomic variations and epigenomic landscape of the Medaka Inbred Kiyosu-Karlsruhe (MIKK) panel. bioRxiv. 2021. p. 2021.05.17.444424. doi:10.1101/2021.05.17.444424

58. Loosli F, Köster RW, Carl M, Kühnlein R, Henrich T, Mücke M, et al. A genetic screen for mutations affecting embryonic development in medaka fish (Oryzias latipes). Mech Dev. 2000;97: 133–139.

59. Abadi M, Barham P, Chen J, Chen Z, Davis A, Dean J, et al. Tensorflow: A system for large-scale machine learning. 12th ${USENIX} symposium on operating systems design and implementation ({OSDI}$ 16). 2016. pp. 265–283.

60. Labelbox: The leading training data platform for data labeling. [cited 6 May 2021]. Available: https://labelbox.com

61. Jung AB, Wada K, Crall J, Tanaka S, Graving J, Yadav S, et al. Imgaug. GitHub: San Francisco, CA, USA. 2020.

62. Head SR, Komori HK, LaMere SA, Whisenant T, Van Nieuwerburgh F, Salomon DR, et al. Library construction for next-generation sequencing: overviews and challenges. Biotechniques. 2014;56: 61–4, 66, 68, passim.

63. Li H, Durbin R. Fast and accurate short read alignment with Burrows-Wheeler transform. Bioinformatics. 2009;25: 1754–1760.

64. McKenna A, Hanna M, Banks E, Sivachenko A, Cibulskis K, Kernytsky A, et al. The Genome Analysis Toolkit: a MapReduce framework for analyzing next-generation DNA sequencing data. Genome Res. 2010;20: 1297–1303.

65. Picard Tools - By Broad Institute. [cited 9 Mar 2020]. Available: http://broadinstitute.github.io/picard/

66. Köster J, Rahmann S. Snakemake--a scalable bioinformatics workflow engine. Bioinformatics. 2012;28: 2520–2522.

67. Leger A. a-slide/NanoSnake: v0.0.3.1. 2020 [cited 17 Feb 2021]. doi:10.5281/zenodo.3630380

68. Chen S, Zhou Y, Chen Y, Gu J. fastp: an ultra-fast all-in-one FASTQ preprocessor. Bioinformatics. 2018;34: i884–i890.

69. Dobin A, Davis CA, Schlesinger F, Drenkow J, Zaleski C, Jha S, et al. STAR: ultrafast universal RNA-seq aligner. Bioinformatics. 2013;29: 15–21.

70. Patro R, Duggal G, Love MI, Irizarry RA, Kingsford C. Salmon provides fast and bias-aware quantification of transcript expression. Nat Methods. 2017;14: 417–419.

71. Casale FP, Rakitsch B, Lippert C, Stegle O. Efficient set tests for the genetic analysis of correlated traits. Nat Methods. 2015;12: 755–758.

72. Purcell S, Neale B, Todd-Brown K, Thomas L, Ferreira MAR, Bender D, et al. PLINK: a tool set for whole-genome association and population-based linkage analyses. Am J Hum Genet. 2007;81: 559–575.

73. Fitzgerald T. ViteRbi. Github; Available: https://github.com/tf2/ViteRbi

74. [No title]. [cited 9 Apr 2021]. Available: ftp://ftp.ensembl.org/pub/release-102/emf/ensembl-compara/multiple_alignments/50_fish.epo/

75. Paradis E, Schliep K. ape 5.0: an environment for modern phylogenetics and evolutionary analyses in R. Bioinformatics. 2019;35: 526–528.

76. R Core Team. R: A Language and Environment for Statistical Computing. Vienna, Austria: R Foundation for Statistical Computing; 2020. Available: https://www.R-project.org/

77. Wickham H, Averick M, Bryan J, Chang W, McGowan LD, François R, et al. Welcome to the Tidyverse. Journal of Open Source Software. 2019;4: 1686.

78. Gu Z, Gu L, Eils R, Schlesner M, Brors B. circlize implements and enhances circular visualization in R. Bioinformatics. 2014;30: 2811–2812.

79. Purcell S, Chang C. PLINK 1.9. [cited 4 Aug 2020]. Available: http://www.cog-genomics.org/plink/1.9/

80. Chang CC, Chow CC, Tellier LC, Vattikuti S, Purcell SM, Lee JJ. Second-generation PLINK: rising to the challenge of larger and richer datasets. Gigascience. 2015;4: 7.

81. R Core Team. R: A Language and Environment for Statistical Computing. Vienna, Austria: R Foundation for Statistical Computing; 2020. Available: https://www.R-project.org/

82. Barrett JC, Fry B, Maller J, Daly MJ. Haploview: analysis and visualization of LD and haplotype maps. Bioinformatics. 2005;21: 263–265.

83. Flynn JM, Hubley R, Goubert C, Rosen J, Clark AG, Feschotte C, et al. RepeatModeler2: automated genomic discovery of transposable element families. Genomics. bioRxiv; 2019. p. 378.

84. Kasahara M, Naruse K, Sasaki S, Nakatani Y, Qu W, Ahsan B, et al. The medaka draft genome and insights into vertebrate genome evolution. Nature. 2007;447: 714–719.

85. Ellinghaus D, Kurtz S, Willhoeft U. LTRharvest, an efficient and flexible software for de novo detection of LTR retrotransposons. BMC Bioinformatics. 2008;9: 18.

86. Ou S, Jiang N. LTR_retriever: A Highly Accurate and Sensitive Program for Identification of Long Terminal Repeat Retrotransposons. Plant Physiol. 2018;176: 1410–1422.

87. Smit AFA, Hubley R, Green P. RepeatMasker home page. 2010. Available: http://www.Repeatmasker.org

88. Inoue Y, Saga T, Aikawa T, Kumagai M, Shimada A, Kawaguchi Y, et al. Complete fusion of a transposon and herpesvirus created the Teratorn mobile element in medaka fish. Nat Commun. 2017;8: 551.

