## Supplementary figures for "The Medaka Inbred Kiyosu-Karlsruhe (MIKK) Panel"

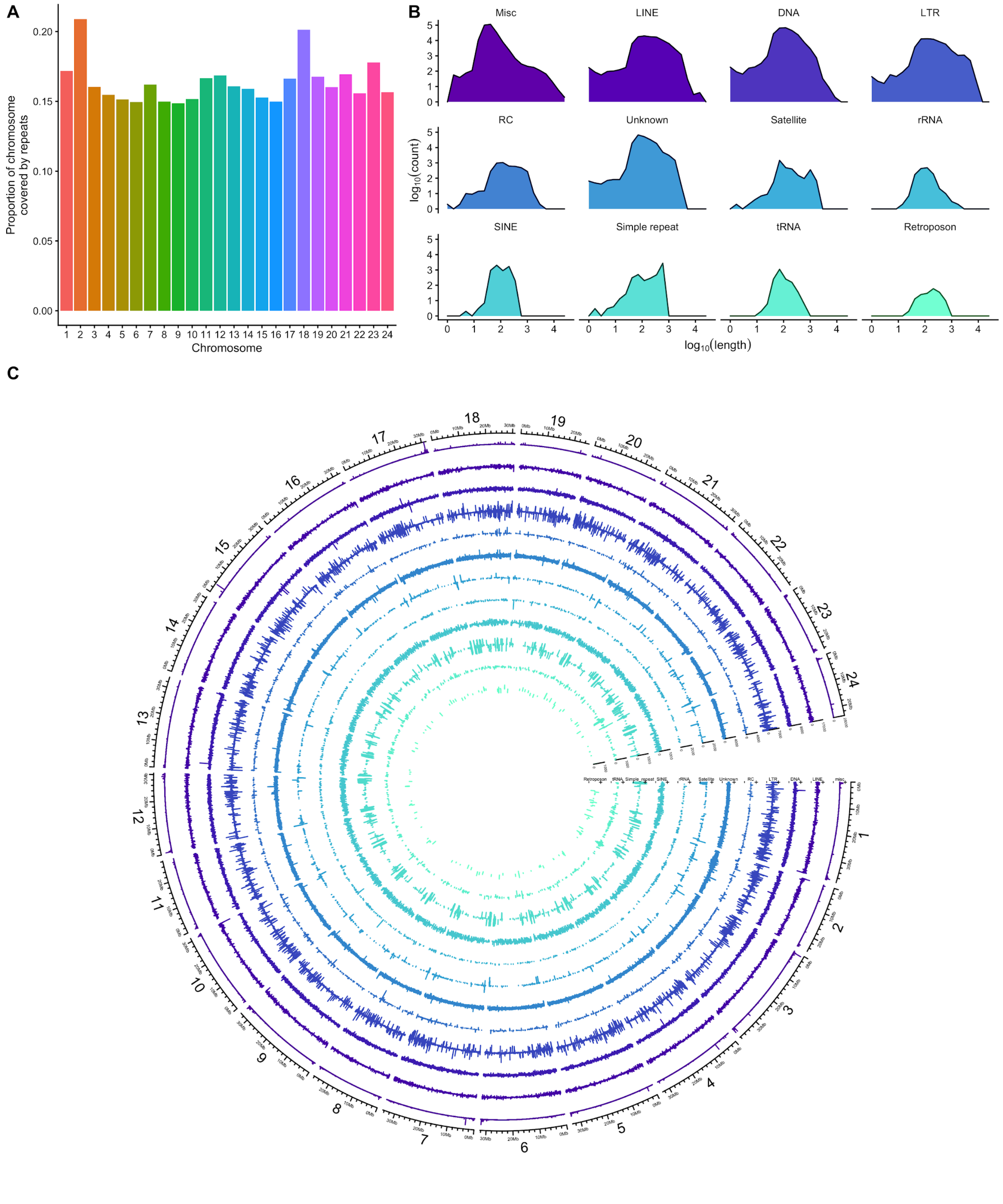


**Supplementary Figure 1:** *Repeat content in the* HdrR *genome based on RepeatMasker results (**Methods**). **A**. Proportion of repeat content per-chromosome. **B**. $log_{10}$ of repeat lengths and counts per repeat class. "Misc" includes all repeats assigned to their own specific class, for example "(GAG)n" or "(GATCCA)n". **C**. Circos plot showing repeat length (radial axes) by locus (angular axis) and repeat class (track). The code and methods used to generate the figure are set out here:* [https://birneylab.github.io/MIKK_genome_main_paper/20210409_repeats.html](https://birneylab.github.io/MIKK_genome_main_paper/20210409_repeats.html)

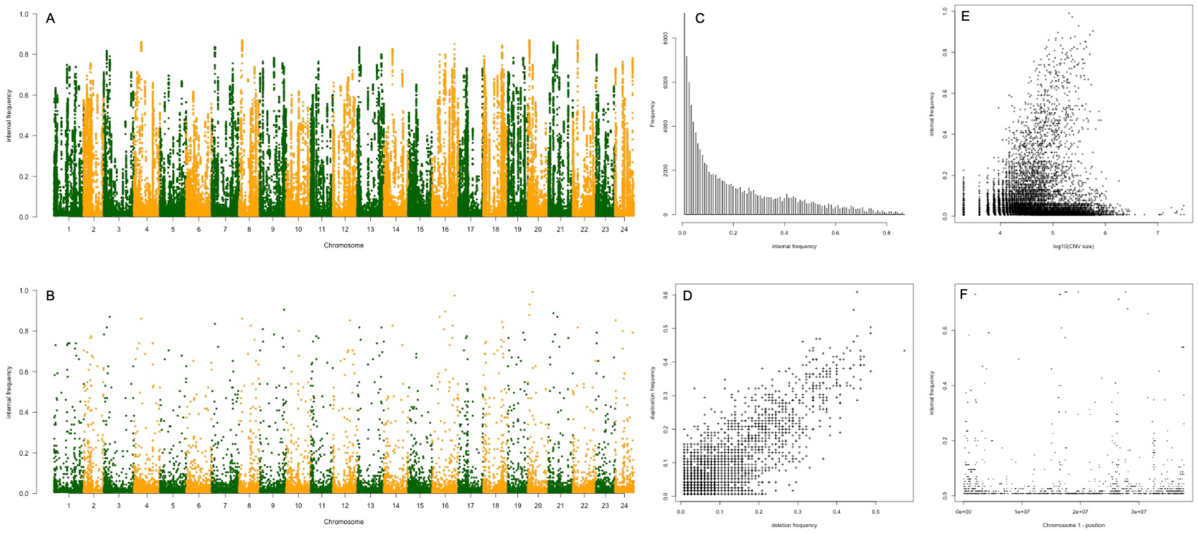


**Supplementary Figure 2*:*** *Assessment of CNV frequency in the MIKK panel. **A**: Overall frequency of all CNV calls across the medaka genome. **B**: Overall combined frequency of merged CNV regions across the medaka genome. **C**: histogram of frequency for all CNV calls. **D**: deletion against Duplication frequency for merged CNV regions. **E**: CNV size against overall frequency for merged CNV regions. **F**: a view showing segments (start and end positions) against overall frequency for all merged CNVs on chromosome 1.*

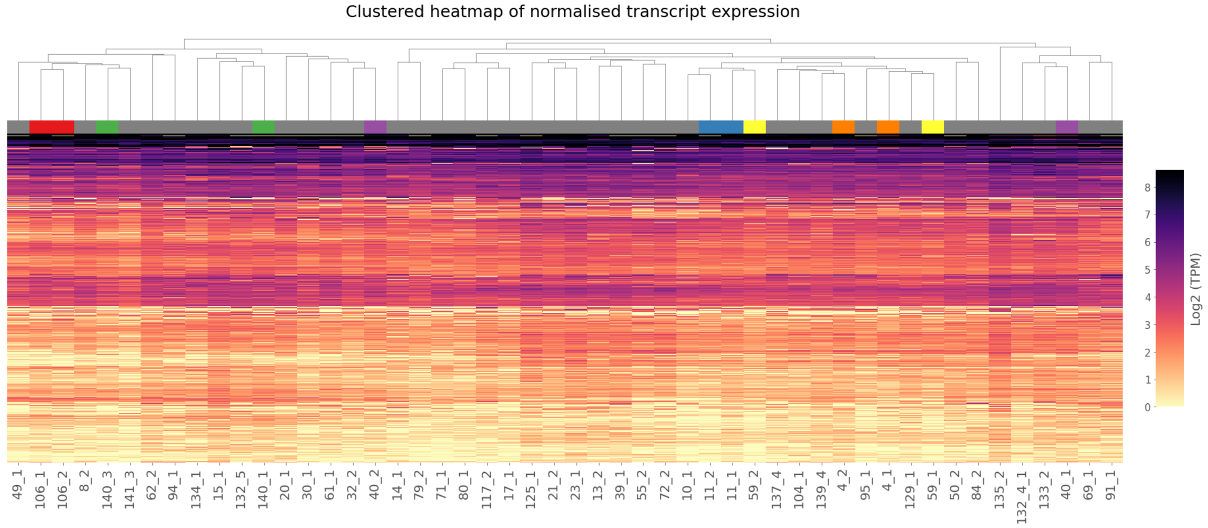


**Supplementary Figure 3:** *Hierarchical clustering of normalised transcript expression across 50 female MIKK panel liver samples, sibling lines are highlighted in different colours.*

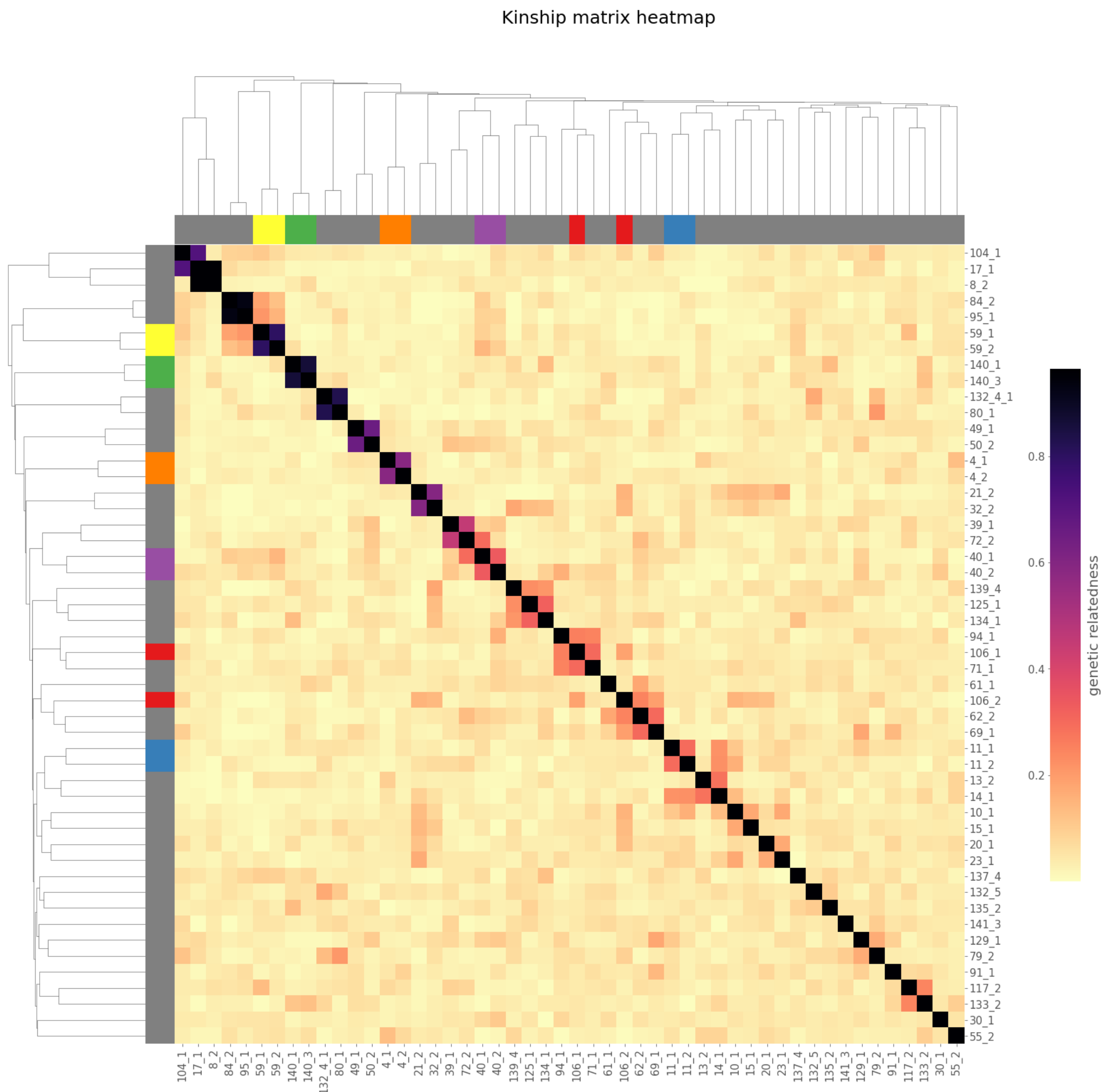


**Supplementary Figure 4:** *Genetic relatedness matrix for the 50 female MIKK panel lines used in the eQTL analysis, derived from SNP genotypes called against the* HdrR *reference. Sibling lines are highlighted in different colours.*
