## Supplementary figures and images for "The Medaka Inbred Kiyosu-Karlsruhe (MIKK) Panel"

### Supplementary file 3

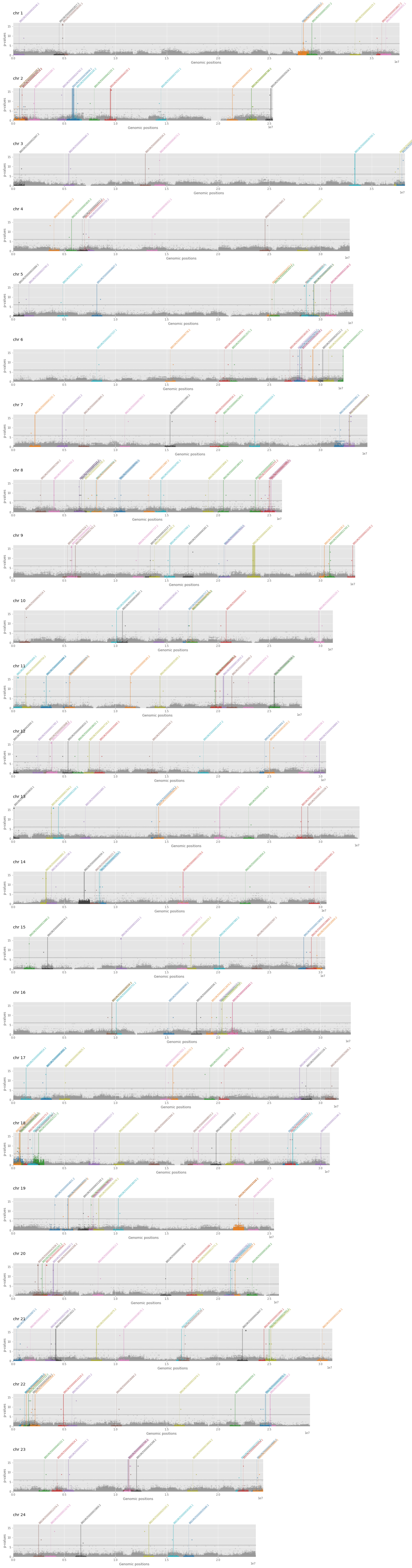

### Supplementary file 4

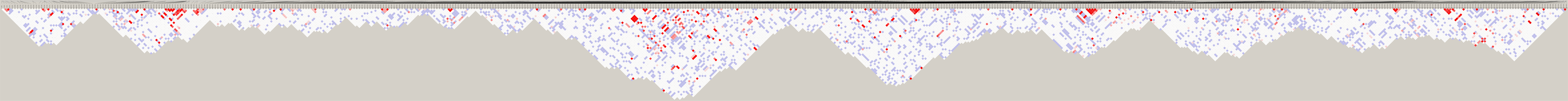

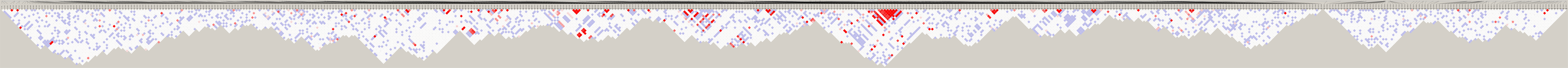

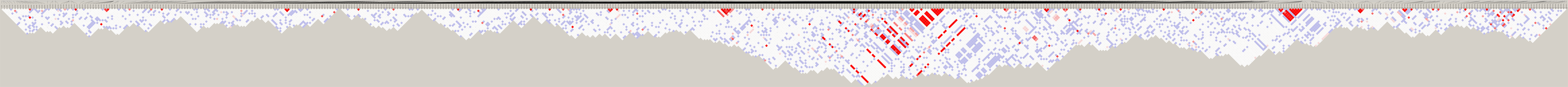
